# Thermodynamic Model and Experiments Indicate Kinetically Trapped States in Ionisable Lipid Nanoparticles

**DOI:** 10.64898/2026.10.28.762589

**Authors:** Ekaterina Kostyurina, Akhil Sudarsan, Marina Huber, Christoph Westerhausen, Nadine Schwierz, Joachim O. Rädler, Susanne Liese

## Abstract

Lipid nanoparticles (LNPs) are the leading delivery vehicles for RNA-based therapies, with pH-dependent mesophase transitions of their ionizable lipids proposed to play a crucial role in LNP function and endosomal escape. However, a consistent quantitative framework linking molecular lipid properties to the thermodynamics of these transitions is still lacking. Here, we establish such a framework by combining X-ray scattering, fluorescence spectroscopy, all-atom molecular dynamics simulations, and continuum modeling. Using small-angle X-ray scattering (SAXS) and Laurdan fluorescence spectroscopy, we characterize the pH- and temperature-dependent phase behavior of both bulk lipid systems and LNPs containing the ionisable lipids MC3 or ALC-0315. Laurdan fluorescence measurements reveal the same pH- and temperature-dependent transitions in bulk and LNP systems, which closely follow the inverse micellar (L_*II*_) – inverse hexagonal (H_*II*_) transitions identified by SAXS, establishing Laurdan as a reporter of these mesophase transitions in LNPs. We combine experimental and simulation data to parameterize a continuum model of the L_*II*_ - H_*II*_ transition, providing a quantitative description of mesophase stability. The model reveals that geometric frustration arising from packing constraints is a key determinant of mesophase stability. By combining experimental and simulation data with theoretical analysis, we determine the relevant membrane parameters, including mean and Gaussian curvature bending rigidities, and identify the parameter ranges that govern phase stability. Together, these results establish a quantitative continuum theory for pH-dependent mesophase transitions of ionisable lipid and suggest metastable states in response to pH changes as an inherent feature of LNP core phase.

## Introduction

Lipid nanoparticles (LNPs) have proven valuable in modern medicine for nucleic acid therapeutics, including mRNA and siRNA.[1] They protect nucleic acids from degradation and facilitate cellular uptake, making them central to many current RNA-based therapies.[2]

Structurally, LNPs are commonly described as core–shell assemblies with a finely tuned composition.[3, 4] The nanoparticle core contains ionizable lipids that complex with nucleic acids and enable encapsulation, cholesterol, which contributes to structural stability, and a hydrated interior surrounding the nucleic acid cargo. The outer shell is stabilized by phospholipids and PEGylated lipids that provide size control and colloidal stability. This multi-component organization is key to both particle stability in blood circulation and functionality during cellular delivery.

During endosomal maturation, the decreasing pH alters the protonation state of ionizable lipids, giving rise to structural transitions within the LNP core (Figure 1a). These mesophase transitions are believed to play an important role in LNP function and have been associated with endosomal escape.[5, 6, 7] Recent studies have associated successful endosomal escape with the ability of ionizable lipids (ILs) in the LNP core to adopt non-lamellar structures, particularly inverted phases such as inverse micellar or inverse hexagonal structures.[5] Inverse hexagonal phases are considered fusogenic and are thought to facilitate membrane destabilization and cargo release.[8, 9] Importantly, SAXS studies on ionizable lipid/cholesterol bulk phases and full LNPs resulted in high resolution insight into the series of mesostructures as a function of pH.[6, 10, 11] In particular, the transition from inverse micellar or inverse hexagonal structures at a critical pH of about 6.0 appeared as a hallmark of various ionizable lipids. As LNPs exhibit signatures of structural transition at the same pH, it is assumed that the transition is critical during endosomal maturation as the endosomal environment becomes progressively more acidic.[6] However, there is no quantitative theoretical models exist that describe how lipid protonation results in structural rearrangement. In his seminal work, Luzzati systematically classified non-bilayer, lipid mesophases and their well-defined thermotropic behavior.[12] These self-assembled structures can be understood in terms of the molecular packing parameter introduced by J. Israelachvili,[13] which relates the effective molecular geometry of amphiphilic lipids—ranging from conical to cylindrical and inverted-conical shapes—to the mesophases they preferentially form.

**Figure 1:**
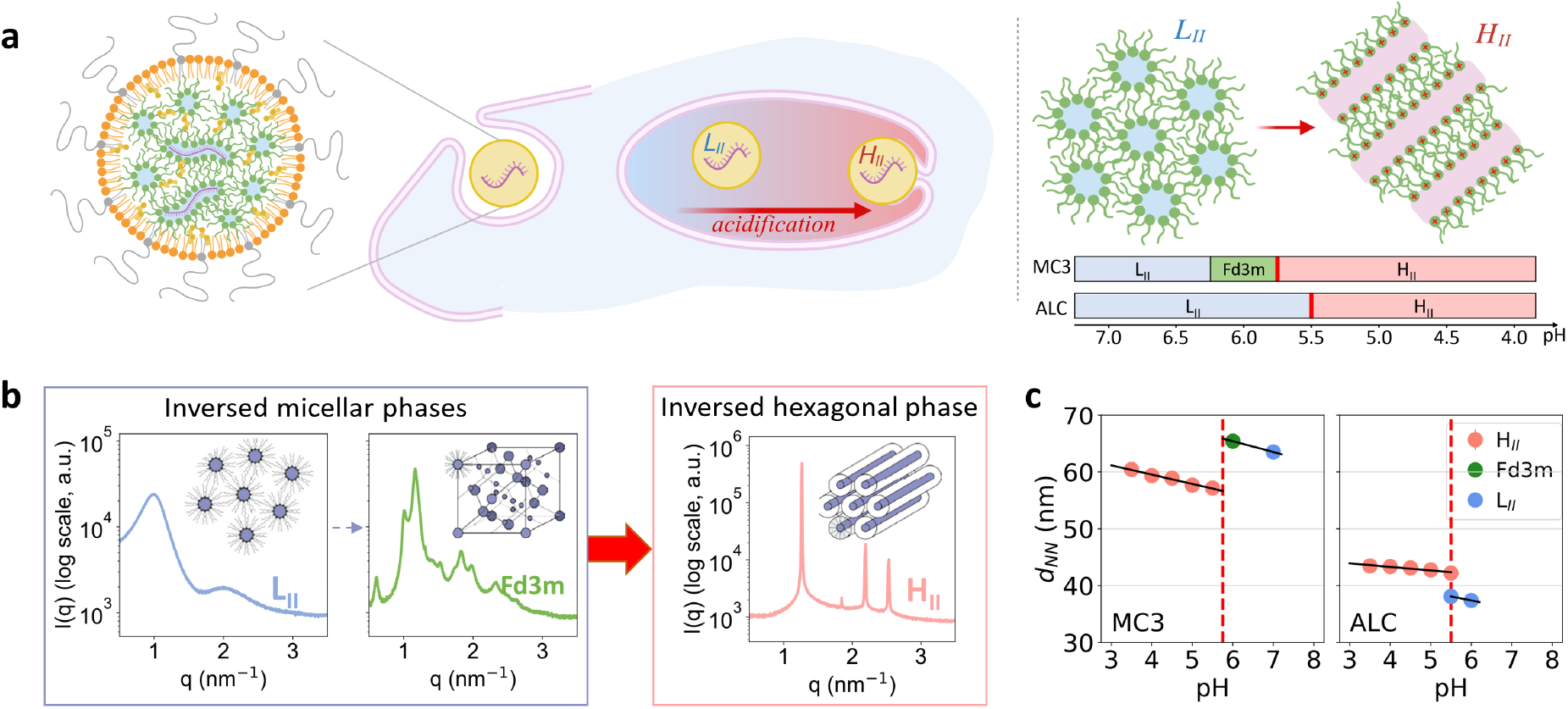
pH-dependent mesophase transitions in LNPs and bulk lipids. **a**.Left: Schematic of an LNP undergoing endosomal uptake and maturation, showing the transition from the L_*II*_ to H_*II*_ core phase during acidification. Right: Cartoon showing the change of internal lipid order corresponding to the L_*II*_–H_*II*_ transition and the phase diagram for MC3/cholesterol and ALC/cholesterol mesophases. **b**. Characteristic SAXS profiles, and the order of their appearance with decreasing pH of MC3/cholesterol as examples of ionizable lipid-cholesterol mesophases, in general. **c**. Next-neighbour distance of MC3/cholesterol and ALC/cholesterol mesophases as a function of pH. The vertical red line shows the transition pH.

To describe the individual lipid phases, the geometry and elastic properties of the lipidic molecular layers is taken into account. In theoretical modeling, this is typically done using the Helfrich energy, which accounts for both the curvature of the lipid interfaces and their bending stiffness.[14, 15] A key aspect of the transition between different inverse lipid phases is the change in topology. During the transition from inverse micellar to inverse hexagonal phases, discrete inverse micelles reorganize into an interconnected array of cylindrical water channels. This transition changes the topology of the lipid-water interface and therefore introduces a significant contribution from the Gaussian curvature term. The Gaussian curvature and the associated bending stiffness are therefore considered a crucial parameter in the description of lipid phases.[16] Additionally, the non-space-filling nature of cylindrical and spherical structures introduces packing frustration.[15, 17, 18] Accordingly, a variety of theoretical models have been developed in the literature to describe the free energy of different inverse mesophases and the transitions between them. While these models differ in their level of detail and underlying assumptions, they share a common framework in which the total free energy consists of a bending contribution, described by the Helfrich elastic energy, and an additional harmonic stretching term that accounts for the molecular rearrangements required to fill space and avoid voids.[19, 20, 18, 21, 22, 15, 23] Most studies have focused on mesophase transitions driven by changes in the water volume fraction or temperature,[19, 20, 21, 15, 17] whereas only a few have considered the influence of lipid tail length or transitions induced by variations in the spontaneous curvature.[18, 22, 23] Many existing models are tailored to specific transitions, particularly within or between bicontinuous and hexagonal phases, while the transition between the inverse micellar L_*II*_ and inverse hexagonal H_*II*_ phases has received comparatively little attention. Furthermore, a consistent application of these concepts to match experimental observations in ionizable lipid bulk phases and LNPs has not yet been performed. A challenge in establishing a quantitative link between theoretical descriptions and experimental measurements lies in the relatively large number of relevant parameters, many of which are difficult to determine independently. Molecular dynamics (MD) simulations help bridge this gap by providing detailed molecular-level insight into the structure and conformational organization of the lipid assemblies. In addition, MD simulations allow the determination of relevant membrane properties, such as the bending rigidity and effective lipid molecular area [24, 25], which can be used to parameterize continuum-level theoretical models. Thus, combining MD simulations with continuum theory provides a means to connect molecular-scale structure and interactions with the mesophase behavior observed experimentally.

Here we present a continuum theory combined with MD and experimental data to explore the free energy landscape of ionizable lipid/cholesterol water phases. We show that elastic energy description of inverse micellar and inverse hexagonal transitions gives rise to a pH-dependent transition. It is found that, in order to meet the experiment, the introduction of packing frustration into the energy contribution is decisive for the transition between different lipid phases. We start by aligning small-angle X-ray scattering (SAXS) and Laurdan fluorescence measurements, which demonstrates that transitions form the L_*II*_ to the H_*II*_ phase occur in LNPs and lipid bulk phases with the same pH and temperature dependence. Subsequently, we develop a theoretical model that identifies packing geometry as a key aspect to facilitated mesophase transitions. Based on fluorescence measurements, we find that LNPs subjected to pH cycling exhibit pronounced hysteresis in the L_*II*_-to-H_*II*_ phase transition, indicating the presence of an energy barrier between these topologically distinct mesophases. The delayed transition upon decreasing pH suggests that the LNP core may transiently persist in a metastable mesophase during endosomal maturation.

## Results and Discussion

### Phase Transitions in IL/Cholesterol Bulk Phases and LNPs

The majority of commonly used ionizable lipids (ILs) exhibit a pH-dependent structural transition from an inverse micellar (L_*II*_) to an inverse hexagonal (H_*II*_) phase within the physiologically relevant endosomal pH range. For the current study, we selected two structurally different ionizable lipids - DLin-MC3-DMA (MC3) and ALC-0315 (ALC)—and studied them in the form of bulk phases, both with and without cholesterol, as well as in LNPs.

First, we characterized the phase behavior and corresponding lattice constants of the bulk phases as a function of pH and, in some cases, temperature using small-angle X-ray scattering (SAXS) (Figure 1b). For both ILs, the L_*II*_–H_*II*_ transition occurs in the pH range of 5–6 (Figure 1a). To compare the characteristic structural dimensions of the different phases, we use the nearest-neighbor distance (*d*_NN_), calculated from the SAXS peak positions. This distance corresponds to the center-to-center distance between neighboring micelles in the L_*II*_ phase or cylinders in the H_*II*_ phase. For both lipids, *d*_NN_ increases with decreasing pH within each phase, with a discontinuous jump around the phase transition (Figure 1c). This trend is consistent with changes in the effective lipid packing parameter resulting from headgroup ionization and with our previous results.[6]

Next we introduce a fluorescence reporter that is capable to detect the L_*II*_–H_*II*_ transition both in ionizable lipid bulk phases as well as in LNPs. Fluorescence spectroscopy allows for faster readout of the state of LNP solutions and can be combined with life cell imaging. However, in order to correctly attribute fluorescence spectra to defined mesophases, samples that were previously characterized using SAXS are required. To observe phase transitions in LNPs, we utilize the solvatochromic fluorescent probe Laurdan. An increase in the hydration level and polarity of its environment enhances dipolar relaxation of the dye, causing a red shift in its emission spectrum and a decrease in the generalized polarization, GP = (I_440_ - I_490_)/(I_440_ + I_490_) (Figure 2a).[26, 27, 28] Laurdan is generally used to determine the phase transition between the ordered gel phase (L_*β*_) and the liquid-crystalline phase (L_*α*_) of lipid membranes. Here, we use it to study the L_*II*_–H_*II*_ transition, which is also associated with significant changes in headgroup hydration upon ionization.[10]

**Figure 2:**
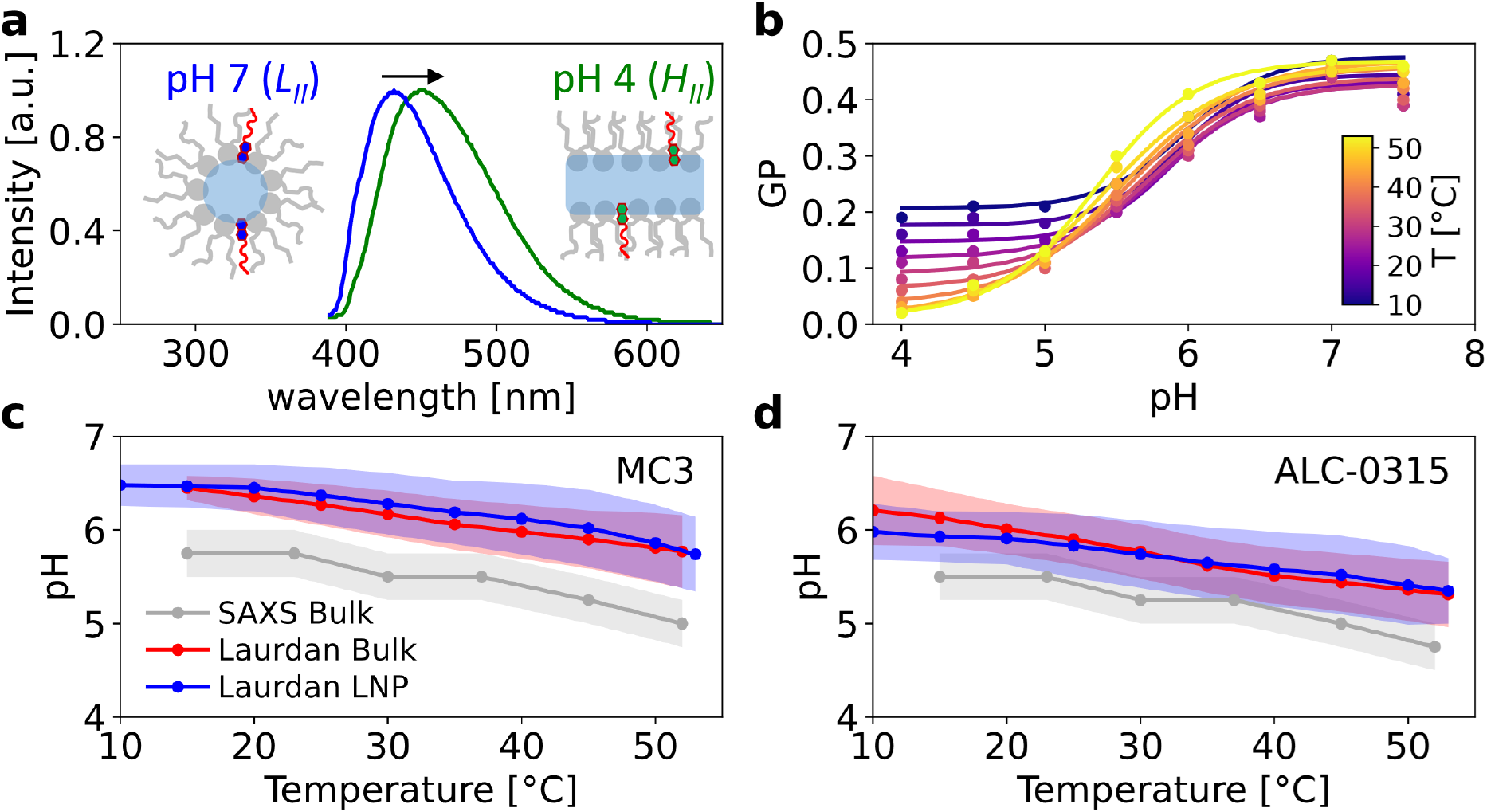
pH- and temperature-dependent phase behavior of bulk systems and LNPs probed by Laurdan and SAXS. **a**.Representative Laurdan emission spectra showing a red shift between the inverse micellar (L_*II*_ , pH 7) and inverse hexagonal (H_*II*_, pH 4) phase of the bulk system (ALC-315, 20 ^*°*^C), resulting in a decreased GP. **b**. Representative pH- and temperature-dependent changes in GP values of the bulk system (ALC-315) indicating a shift of the L_*II*_ – H_*II*_ transition toward lower pH* values as the temperature increases. Values of the transition pH* determined from the inflection points of the Boltzmann sigmoid fits. **c**. Temperature-dependent transition pH* of the bulk system and LNPs formulated with DLin-MC3-DMA determined using the above-mentioned Laurdan-based method and compared with values obtained by SAXS. **d**. Temperature-dependent transition pH* of the bulk system and LNPs formulated with ALC-315 determined using the above-mentioned Laurdan-based method and compared with values obtained by SAXS.

To demonstrate the consistency between SAXS and fluorescence spectroscopy, we first measured the GP of Laurdan-containing bulk phases as a function of pH and temperature. The transition pH values (pH*) determined from the inflection points of Boltzmann sigmoid fits (Figure 2b) are in good agreement with those obtained from SAXS (Figure 2c,d). With both methods we measure that the transition pH decreases with increasing temperature following the same trend. The small shift in pH* may arise from the different definitions of the phase transition in the two methods. Whereas SAXS measures the lipid phases directly, GP reflects changes in headgroup solvation, which may follow slightly different behavior. Indeed, when the pH* obtained from SAXS is mapped onto the GP(pH) curve, it is located at the point where GP begins to increase (Figure 2b). However, as both methods show the same temperature dependence of pH*, the transitions detected by the two approaches most likely have the same underlying origin.

As a next step, we performed the same measurements on LNPs (Figure 2c,d, red data). The observed trends follow those of the bulk phases remarkably closely, supporting the conclusion that the thermodynamics we study in bulk phases also govern the corresponding transitions occurring within LNPs.

### Continuum Model of Mesophase Transitions

To rationalize the L_*II*_–H_*II*_ transition, we develop a continuum description based on interfacial curvature elasticity. A comparison of the energies of the L_*II*_ and H_*II*_ phases shows that curvature contributions alone are insufficient to account for their stability and the observed topology change. We therefore extend the model in a second step to include additional contributions arising from lipid packing constraints.

The lipid–water interface is described within the standard Helfrich framework for curvature elasticity

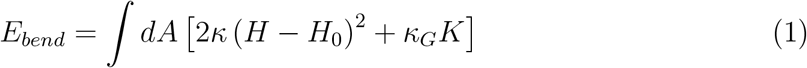

where the integral is taken over the interfacial area *A, H* and *K* denote the mean and Gaussian curvatures, *κ* and *κ*_*G*_ are the corresponding bending rigidities, and *H*_0_ is the spontaneous curvature.

The total number of lipids is conserved. Moreover, lipid interfaces allow only very limited stretching, such that the total area *A*_*tot*_ remains constant at fixed pH or temperature. Terms linear in area therefore contribute only a constant offset and do not affect energy minimization. At a given pH and temperature, we further assume that a spontaneous curvature is established, characterized by a curvature radius 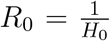. Increasing pH is associated with deprotonation and a smaller spontaneous curvature radius.

The H_*II*_ and L_*II*_ phases are modeled as assemblies of *N*_*S*_ spheres with radius *R*_*S*_ and *N*_*C*_ cylinders with radius *R*_*C*_ and length *L*, respectively. The radius *R*_*S*_ and *R*_*C*_ set the mean and Gaussian curvature of the respective shape (Figure 3a). The corresponding bending energies read

**Figure 3:**
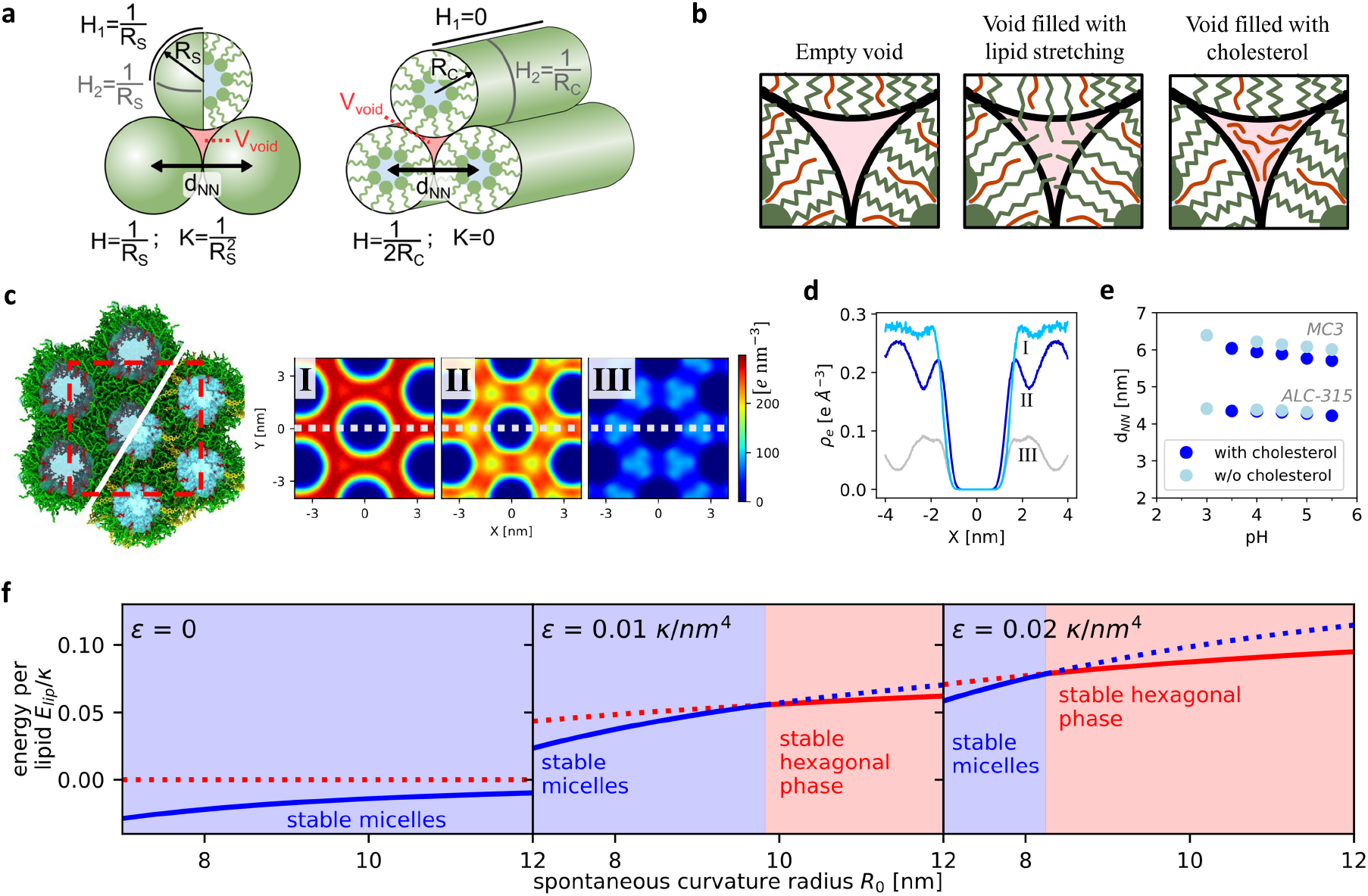
Packing Constraint and Void Filling **a.**Schematic illustration of the characteristic curvatures of spherical and cylindrical shapes. **b**. Schematic illustration of empty and filled void volume. **c**. Snapshot of MD simulation of MC3 lipid (upper-left half) and MC3/cholesterol mixture (lower-right half) in HII phase, and 2D electron-density maps of MC3 lipid tails in MC3-only system (I), MC3 lipid tails in MC3/cholesterol system (II), and cholesterol in MC3/cholesterol system (III), obtained from MD simulation. **d**. Electron-density distribution cut along the marked axes of 2D electron-density maps in figure c (I, II, III). **e**. Next-neighbor distances of MC3 and ALC-315 lipids with and with-out cholesterol obtained from SAXS. **f**. Exemplary energy curves for different 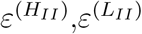. To keep the examples minimal, we set 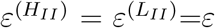. The energy of the stable phase is shown as a solid line, while the energy of the unstable phase is shown as a dotted line. A finite value of 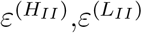 is required to observe a mesophase transition.

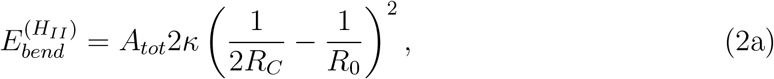

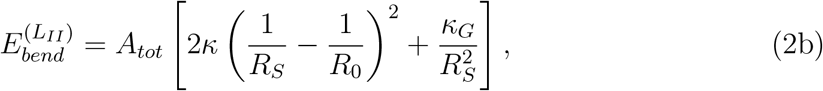

where the total interfacial area is *A*_*tot*_ = 2*πR*_*C*_*LN*_*c*_ in the H_*II*_ phase and 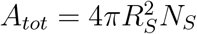 in the L_*III*_ phase. The bending energy is minimized with respect to *R*_*S*_ and *R*_*C*_ at fixed *A*_*tot*_, while the volume is allowed to adjust through water redistribution. This yields 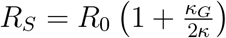 and 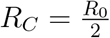 , and thus

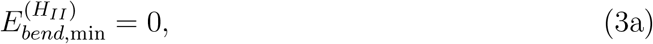

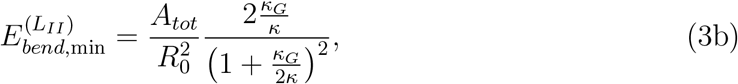

For wide range of lipid systems, the ratio 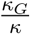 is negative.[29] This would imply that the L_*II*_ phase is always energetically favoured, in clear contradiction to experimental observations.

It is worth noting that phase transitions between planar, cylindrical, and spherical lipid interfaces have been successfully described in alternative theoretical approaches. These models typically impose additional constraints, such as a fixed surface-to-volume ratio, which effectively limits the accessible water content.[30, 14] In contrast, the present model allows the volume to equilibrate freely, corresponding to systems in which water redistribution is not kinetically hindered. Under these conditions, bending energy alone is insufficient to reproduce the experimentally observed mesophase behavior, indicating that additional energetic contributions must be taken into account.

### Void Filling and Effect of Cholesterol

An additional energetic contribution arises from the incompatibility between maintaining perfectly spherical or cylindrical molecular geometries and filling space without leaving voids. This geometric mismatch has been described in the literature using terms such as packing frustration, space filling cages, and bending frustration.[18, 31, 15, 17, 23] To eliminate voids within the lipid assembly, molecular rearrangements must occur in the interfacial region, allowing the structure to accommodate the constraints imposed by efficient space filling.

Molecular dynamics simulations of the hexagonal phase, for both MC3- and ALC-based systems with and without cholesterol, confirm the absence of empty voids between adjacent cylindrical assemblies (Figure 3c). Electron density maps show that, in the absence of cholesterol, these void regions are filled by lipid tails, whereas in cholesterol-containing systems cholesterol preferentially accumulates in the interstitial regions between neighboring cylinders, which can be quantitatively observed in the horizontal cuts in Figure 3d. These observations suggest that, as a helper lipid, cholesterol reduces the packing penalty by occupying interstitial regions and thereby alleviating the need for lipid tail deformation in highly curved mesophases. Consistent with this molecular picture, SAXS measurements show that cholesterol condenses the inverted phases by reducing the nearest-neighbor distance of the cylinders. This effect is pronounced for the less strongly curved MC3 system but is much weaker for the more highly curved ALC (Figure 3e), indicating that the influence of cholesterol depends on the intrinsic curvature of the ionizable lipid. Together, these observations provide the molecular basis for the continuum description developed below.

In cholesterol-free systems, void filling is dominated by lipid tail stretching and packing distortions, (Figure 3b). In IL/cholesterol systems the energy contributions associated with void filling arise from several microscopic mechanisms, including lipid tail stretching, the packing factor, and cholesterol redistribution. A full molecular description of the energetics of inverse lipid mesophases is inherently complex. As discussed in the Introduction, theoretical descriptions of inverse mesophases commonly combine the Helfrich bending energy with a harmonic stretching contribution to account for packing frustration and the molecular rearrangements required for space filling.[19, 20, 18, 21, 22, 15, 23] Since void filling is governed by multiple competing mechanisms that cannot be fully captured by existing models, we do not directly adopt a specific literature model, but instead develop a scaling-based description. Our approach captures the essential geometric dependence of the relevant energetic contributions and is suitable for describing the mesophase transitions considered here.

We note that the number of molecules involved to form a spherical or cylindrical structure scales with the total interfacial area *A*_*tot*_, while the local deformation or compositional mismatch that must be accommodated scales with the characteristic radius *R*. Assuming a harmonic energetic penalty for these deviations, both lipid stretching and cholesterol redistribution lead to the same scaling behaviour,

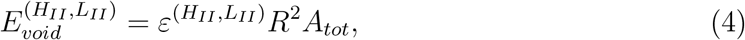

where 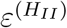 and 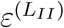 are phenomenological prefactor that sets the magnitude of the void-filling energy. The total energy now reads:

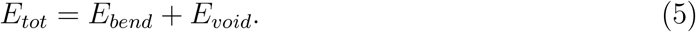

To illustrate the impact of the void energy term, Fig. 3f shows the energy per lipid in both mesophases for various 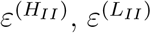. To keep the example concise, 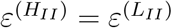 is set. Additionally, values of *κ*_*G*_*/κ* = *−*1 and the area per lipid *a*_*lip*_ = 0.7 nm^2^ are assumed. We note that the exact values of *κ*_*G*_*/κ* and *a*_*lip*_ influence the shape of the curve quantitatively but not qualitatively. As expected, for 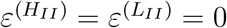, there is no region with a stable hexagonal phase. As 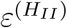 and 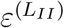 increase, the transition between the mesophases shifts toward smaller values of the spontaneous curvature radius, that is, higher pH values.

### Combining theory, experiment and MD for quantitative description of ionizable lipids

The theory described above allows for a quantitative prediction of ionizable lipid/cholesterol polymorphism. Such an analysis, however, requires a comprehensive set of data, as no single method can determine all relevant parameters. This includes experimental measurements of the pH-dependent phase behavior and nearest-neighbor distances as a function of osmotic pressure, together with molecular dynamics simulations and fitting of the continuum model parameters. To keep this integrated analysis tractable and internally consistent, we focus on MC3 and ALC lipid systems, for which all required datasets can be combined within a common framework.

Equations 2a and 2b contain seven model parameters:: *R*_*C*_, *R*_*S*_, *R*_0_, *κ, κ*_*G*_, 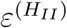, 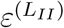. The mean curvature radii *R*_*c*_ and *R*_*s*_ are derived from the experimentally measured nearest-neighbour distances *d*_*NN*_ and position of the pivotal plane determined from MD as described in the next section. The bending modulus (*κ*) is calculated from the splay distribution in MD simulation (see next section). This leaves four parameters to be determined. At the mesophase transition, three additional constraints follow from the equilibrium conditions of the two phases. Firstly, for each phase, the observed equilibrium geometry corresponds to a minimum of the free energy with respect to its curvature radius, giving the conditions

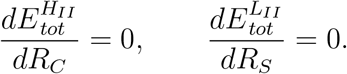

Secondly, assuming thermodynamic equilibrium at the transition pH (pH*), the free energies of the two phases are equal:

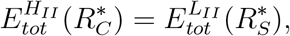

where 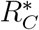 and 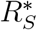 denote the mean curvature radii of cylinder and sphere at pH*.

However, these three conditions are insufficient to determine the four remaining parameters. We therefore obtain an additional constraint by measuring *d*_*NN*_ as a function of osmotic pressure and fitting the resulting dependence as described further. Together, these complementary measurements allow us to determine all seven quantities, with the mean- and spontaneous-curvature radii being a function of pH. In the next two sections, we describe the MD simulations and osmotic-pressure experiments in more detail.

### Bending rigidity of ionizable lipid mesophases from molecular simulations

To establish a quantitative framework for the mesophase transitions of ionizable lipids, the elastic properties governing the energetic cost of membrane deformation need to be determined. A central parameter is the bending rigidity (*κ*), which sets the energetic penalty for deviations from the preferred monolayer curvature and therefore contributes directly to the relative stability of different mesophases. We determined the bending rigidity of the ionizable lipid mixtures directly from all-atom MD simulations of the H_*II*_ phases (Figure 4a,b).

**Figure 4:**
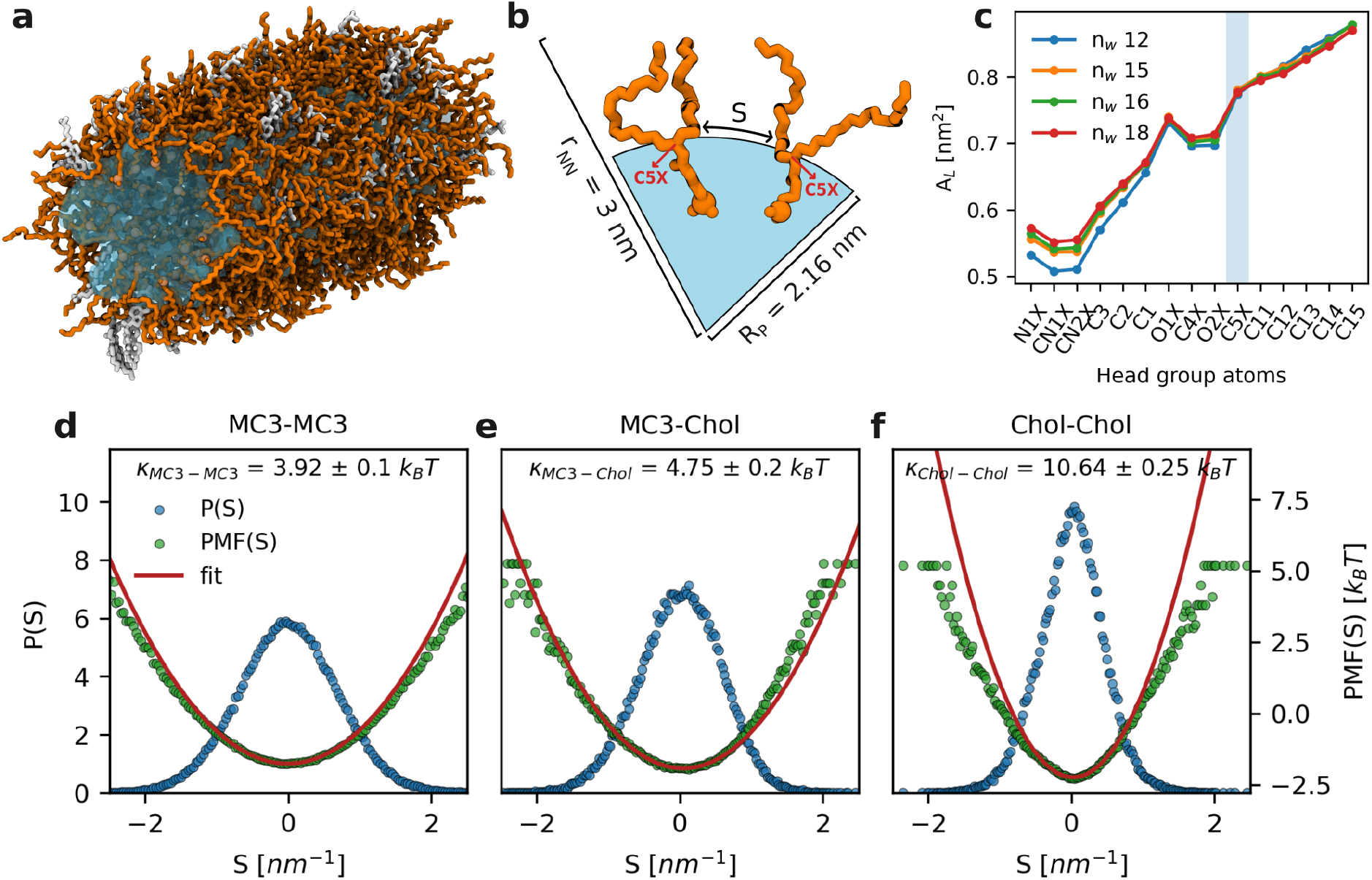
Derivation of the bending rigidity of ionizable lipids from MD simulations **a.**Representative simulation snapshot of the MC3/cholesterol H_*II*_ phase. MC3, cholesterol and water are represented in orange, white and blue colors respectively **b**. Schematic showing the definition of lipid splay (S) between two adjacent lipids. Here r_*NN*_ denotes half the lattice spacing d_*NN*_ and R_*P*_ represents the pivotal plane measured from the center of the H_*II*_ phase. **c**. The area per lipid obtained at different positions corresponding to each atom is shown for MC3 based H_*II*_ phases with different water contents. n_*w*_ represents the number of water molecules per lipid. The pivotal plane is defined at the position of the C5X atom. Lipid splay distributions (blue), the corresponding potentials of mean force (green) and the parabolic fits to the PMFs (red) obtained for **d**. MC3-MC3 **e**. MC3-Chol and **f**. Chol-Chol lipid pairs with their corresponding bending rigidities obtained from MD simulations of MC3/Cholesterol H_*II*_ phase. The effective bending rigidity was then obtained by a pair-count weighted average described in Section S5.1.

Relating molecular fluctuations to continuum curvature elasticity requires the identification of an appropriate reference surface within the lipid layer. Across H_*II*_ phases with different hydration levels and thus different curvatures, the surface area varied along the lipid atoms but became approximately independent of curvature near the junction between the head group and hydrocarbon tails (Figure 4c). For MC3, this pivotal plane was localized at the position of the C5X atom (Figure S9b) . This location is consistent with previous studies of conventional phospholipids, where the pivotal plane has similarly been localized near the headgroup–tail junction [24, 25]. An equivalent position was assumed for ALC H_*II*_ phases.

At the pivotal plane, the probability distributions of local lipid splay were approximately Gaussian for IL–IL, IL–cholesterol, and cholesterol–cholesterol pairs (Figure 4d–f for MC3 and Figure S10 for ALC). The corresponding potentials of mean force showed a quadratic dependence in the low-splay regime, consistent with the harmonic approximation underlying continuum elastic theory. The curvature of these potentials provides the corresponding pairwise contributions to the bending rigidity following the real-space splay approach of Khelashvili and co-workers. The resulting bending rigidities for the different H_*II*_ phases are summarized in Table 1 and provide the molecularly derived elastic parameters for the continuum model of mesophase stability developed below.

**Table 1:** Resulting set of model parameters obtained from the joint fit of experimental/MD data. Comparison of MC3 and ALC lipids with and without cholesterol. The transition pH* is obtianed from SAXS (blue), bending rigidity (***κ***) from MD (orange), Gaussian bending rigidity (***κ***_**G**_), spontaneous curvature radius (**R**_**0**_) and void energy prefactors 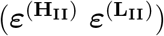– from joint fit by modeling (Green)

| | | pH* | $\kappa$, $kT$ | $\kappa_G$, $kT$ | $R_0$(pH*), nm | $\varepsilon^{(HII)}$, $kT\text{ nm}^{-4}$ | $\varepsilon^{(LII)}$, $kT\text{ nm}^{-4}$ |
| --- | --- | --- | --- | --- | --- | --- | --- |
| MC3 | w/o chol. | 5.75 | $3.78 \pm 0.04$ | $-4.4 \pm 0.5$ | $13 \pm 30$ | $0.15 \pm 0.04$ | $0.14 \pm 0.02$ |
| | with chol. | 5.75 | $4.39 \pm 0.1$ | $-4.2 \pm 0.2$ | $8.5 \pm 2.0$ | $0.24 \pm 0.03$ | $0.173 \pm 0.009$ |
| ALC-315 | w/o chol | 5.5 | $4.1 \pm 0.24$ | $-3.70 \pm 0.09$ | $2.28 \pm 0.06$ | $0.02 \pm 0.04$ | $1.15 \pm 0.02$ |
| | with chol. | 5.5 | $2.7 \pm 0.1$ | $-2.64 \pm 0.02$ | $1.97 \pm 0.02$ | $0.01 \pm 0.02$ | $1.84 \pm 0.05$ |

### Varying osmotic pressure

The experimental *d*_*NN*_ (*p*) curves have been obtained from SAXS. The pressure has been applied to the lipid bulk phases by emerging them into PEG-8k solutions of concentration 15 - 50 wt% resulting in osmotic pressures ranging from 0.4 to 10 MPa (Figure 5a).[32] As the polymer size is significantly larger than the size of the water pores in the lipid phases, water has to be pushed out of the phases to equilibrate the pressure, leading to the decrease in *d*_*NN*_ observed by SAXS (Figure 5b,c).

**Figure 5:**
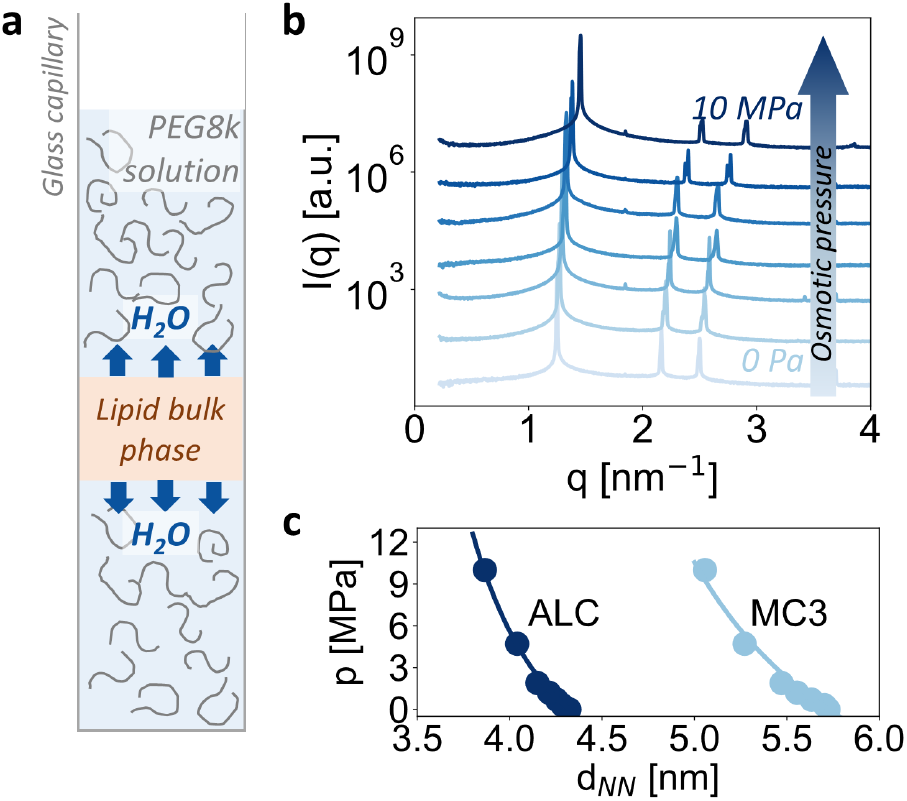
Osmotic pressure experiments. **a.**Schematic illustration of the osmotic pressure setup. Varying PEG concentration in solution allows to apply desired osmotic pressure to a lipid bulk phase, which respond by expelling water from the phase. **b**. Exemplary SAXS curves for MC3/Cholesterol mesophases at pH 5. **c**. Next-neighbour distance as a function of osmotic pressure for MC3/cholesterol mesophases at pH 5.5, and ALC/cholesterol mesophases at pH 5.0. Solid lines show the best fit with the theoretical model (eq. 7).

The applied pressure introduces an additional energetic contribution that scales with the volume of the water domains, such that the total free energy is given by

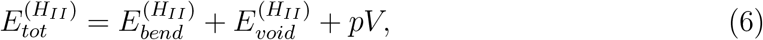

with the volume 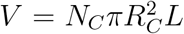. Using Eqs. 2 and 4 and minimizing with respect to *R*_*C*_ leads to

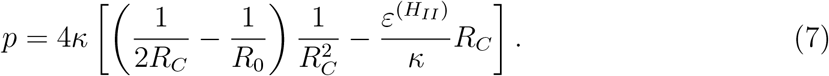

Using MD-derived parameters and the experimentally determined change in next-neighbor distance at the transition point, Eq. 7 can be reduced to a function of a single unknown parameter, the Gaussian bending rigidity *κ*_*G*_, which is used as the fitting parameter. The best fits for MC3/cholesterol and ALC/cholesterol lipid systems are shown in Figure 5c and the details of the fitting procedure are provided in the Supporting information.

## Discussion of the fit results

The analysis described above was performed for two structurally distinct ionizable lipids, MC3 and ALC, both with and without cholesterol. By combining SAXS, MD simulations, and continuum modeling, we determined the transition pH, the pH-dependent spontaneous curvature radius, mean and Gaussian bending rigidities, and the void-filling energy factors of the L_*II*_ and H_*II*_ phases. Together, these parameters allow us to reconstruct the lipid free-energy landscape as a function of spontaneous curvature and, consequently, pH (Figure 6). The resulting parameters are summarized in Table 1. Despite substantial differences between the two lipids, both systems can be described within the same theoretical framework, indicating that their phase behavior is governed by the same underlying thermodynamic principles. Notably, the model also captures the opposite directions of the *d*_*NN*_ discontinuity observed for MC3 and ALC at the phase transition: *d*_*NN*_ decreases for MC3 but increases for ALC upon decreasing pH. We note that the curvature radius *R* also exhibits a jump analog to the discontinuous transition in *d*_*NN*_ , since the next neighbour distance and the curvature radius are connected via *R* = *d*_*NN*_ */*2 *− ι*, with the offset between the next-neighbor distance and the pivotal. From MD simulations we find *ι* = (1.0 *±* 0.3) nm for MC3 and (1.0 *±* 0.4) nm for ALC.

**Figure 6:**
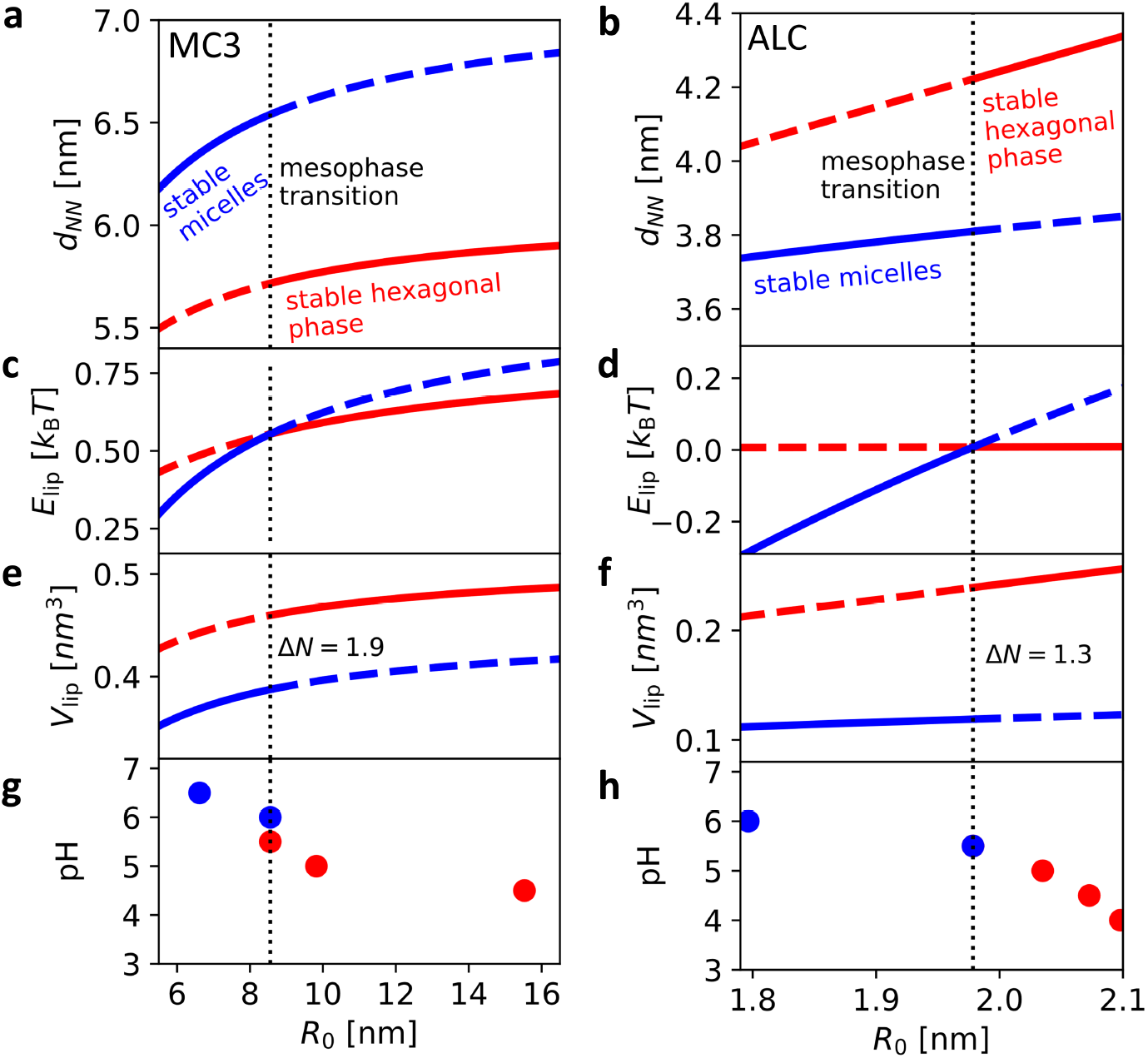
Numerical calculation of lipid free energy and volume per lipid as a function of spontaneous curvature and pH. The curves are obtained using the fitted parameters (Table 1) for the phase transition for MC3/cholesterol (left column) and ALC/cholesterol (right column) systems. **a**,**b**.Fitted dependence of next-neighbour distance (d_*NN*_) on the spontaneous curvature radius (R_0_), **c**,**d**. Calculated energy per lipid (E_*lip*_) as a function of R_0_. At the transition the average number of water molecules per lipids has to increase by Δ*N* , **e**,**f**. Calculated volume per lipid (V_*lip*_) as a function of R_0_, **g**,**h**. R_0_ as a function of pH obtained using experimental d_*NN*_ (pH) data (ref to Fig.1c).

For both lipids, the total system volume increases upon the transition from L_II_ to H_II_ (Figure 6, see Supporting Information for the detailed derivation of *V*_lip_). To compare the two phases independently of system size, we normalize the total volume by the total number of lipids and refer to this quantity as the volume per lipid. For ALC, the increase in volume per lipid may appear counterintuitive because the curvature radius decreases at the transition. However, the volume is not determined by the curvature radius alone, but also by the geometry of the structure. For a fixed number of lipids, and therefore a fixed surface area, a spherical structure encloses less volume than a cylindrical one at the same characteristic radius. Conversely, when comparing the L_II_ and H_II_ geometries, the transition to the more highly curved H_II_ structure can therefore be accompanied by an increase in enclosed volume despite the smaller radius. The resulting increase in volume requires additional water uptake, corresponding on average to Δ*N* = 1.9 water molecules per lipid for MC3 and Δ*N* = 1.3 for ALC. In LNPs, this change in the lipid-to-water ratio can therefore be accommodated either by taking up water or, if water uptake is not possible, by reducing the number of lipids in the system. The latter would require substantial lipid rearrangement to accommodate the increased volume and could thereby contribute to the release mechanism.

Changes in volume per lipid also affect the void volume. Assuming closest packing, spheres have a packing fraction of 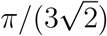, whereas parallel cylinders have a packing fraction of 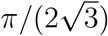. Thus, the void volume per lipid is approximately 0.35 *V*_lip_ in micellar] and 0.10 *V*_lip_ in hexagonal structures. Consequently, the transition from L_II_ to H_II_ reduces the available void volume (see Figure S3), requiring excess cholesterol to be expelled.

The most pronounced difference between the two lipids lies in their spontaneous curvature radius, *R*_0_. ALC exhibits a substantially smaller *R*_0_, corresponding to a higher preferred curvature, consistent with its highly branched molecular structure.

Interestingly, the bending rigidities of the two lipids are similar in the absence of cholesterol. Cholesterol, however, affects the two systems in opposite ways, increasing the stiffness of MC3 mesophases while decreasing that of ALC. Such a lipid-dependent response is consistent with previous observations that the effect of cholesterol on membrane elasticity depends strongly on lipid tail architecture.[33] Thus, rather than acting as a universal membrane stiffener, cholesterol modulates membrane elasticity in an architecture-dependent manner, with consequences for the energy landscape inside LNPs. This observation is particularly interesting in light of the growing number of studies exploring cholesterol modifications as a strategy for modulating LNP biodistribution and cell-specific targeting.[34, 35, 36]

The Gaussian bending rigidity, *κ*_*G*_, is another key parameter contributing to topological phase transitions and membrane fusion. A more negative *κ*_*G*_ stabilizes the positive Gaussian curvature of inverse micelles while increasing the energetic cost of saddle-like structures with negative Gaussian curvature. For all studied systems, the ratio *κ*_*G*_*/κ* is close to *−*1, which is typical for lipid membranes. Nevertheless, the magnitude of *κ*_*G*_ differs between the two lipids. In the presence of cholesterol, *κ*_*G*_ of MC3 is about one and a half times as large as that of ALC, which implies a higher energetic cost of forming saddle-like structures essential for fusion process. In an approximation of two fusing vesicles the cost associated with Gaussian curvature change reads as Δ*G*_*fusion*_ = *−* 4*πκ*_*G*_,predicting higher fusogenity of ALC lipid as compared to MC3.

It is also informative to compare the void-energy prefactors of the hexagonal and mi-cellar phases, 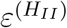 and 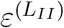, which characterize the energetic cost of void filling in the two geometries. For all MC3 systems, the two prefactors are of comparable magnitude. In contrast, for the highly branched ALC lipid, 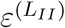 exceeds 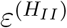 by nearly two orders of magnitude, both with and without cholesterol. This pronounced difference indicates that void filling is energetically much more costly in the micellar than in the hexagonal geometry for ALC. On the other hand, the molecular packing argument of Israelachvili would predict highly curved inverse micellar phases for ALC, in line with its large spontaneous curvature. The opposing tendencies of spontaneous curvature and void-filling energy emphasize that mesophase stability cannot be understood from molecular packing or curvature elasticity alone, but results from a balance between these contributions. The pronounced effect of lipid tail architecture on the constraints associated with void filling is, however, not intuitively evident from the molecular structure alone and points to the need for a deeper understanding of lipid tail dynamics, particularly for the complex architectures of modern synthetic ionizable lipids.

## Metastable States in Lipid Nanoparticles

The analysis above provides an equilibrium description of the L_*II*_–H_*II*_ transition and its associated free-energy landscape. However, the experimentally observed transition may additionally be influenced by metastability and kinetic barriers. To probe these nonequilibrium effects, we monitored the mesophase transition in LNPs using Laurdan fluorescence while cycling the pH from 8 to 4 and back. Both lipids exhibited clear hysteresis, with ΔpH=0.30 for MC3-containing LNPs and ΔpH=0.13 for ALC-containing LNPs (Figure 7a,c). This hysteresis indicates that the LNPs can remain in a metastable mesophase beyond the equilibrium transition pH. A possible microscopic picture of the transition is that the LNP core does not transform uniformly, but instead undergoes a local structural rearrangement that subsequently propagates through the particle. At high pH, the LNP core is predominantly in the L_II_ phase. Upon decreasing the pH, the transition may be initiated locally by the conversion of a small region from the L_II_ to the H_II_ structure. We denote the energy difference between the pure L_II_ state and this intermediate transition structure as Δ*E*_trans_. This local rearrangement establishes the energetic cost associated with initiating the switch between the two mesophases and can trigger the subsequent transformation of the remaining LNP core. At sufficiently low pH, the transformation is complete and the entire LNP core adopts the H_II_ structure. A similar mechanism can be envisioned for the reverse transition upon increasing the pH, with a local region of the H_II_ phase first rearranging toward the L_II_ structure, followed by propagation of the transformation throughout the particle. Within this picture, the observed transition can be approximated as an effective three-state process. Using the fluorescence curves and a three-state model, we estimate Δ*E*_trans_ to be on the order of 1 *k*_B_*T* (see SI for a detailed derivation). Furthermore, the results obtained in Fig. 6 provide the energy differences per lipid for varying pH (Fig.7b,e). Together with the experimentally observed shift ΔpH we estimate an energy difference on the order of magnitude of 0.01 *k*_B_*T* per lipid at the transition (Fig.7c,f). Comparing this per-lipid energy scale with the free-energy offset per LNP indicates that approximately 100 lipids must participate in the initial cooperative rearrangement to initiate the transition. Since an individual LNP contains several thousand lipids, the free-energy change associated with complete conversion of the particle can readily exceed 100 *k*_B_*T* . Such a collective energy release may contribute to membrane destabilization and facilitate endosomal escape.

**Figure 7:**
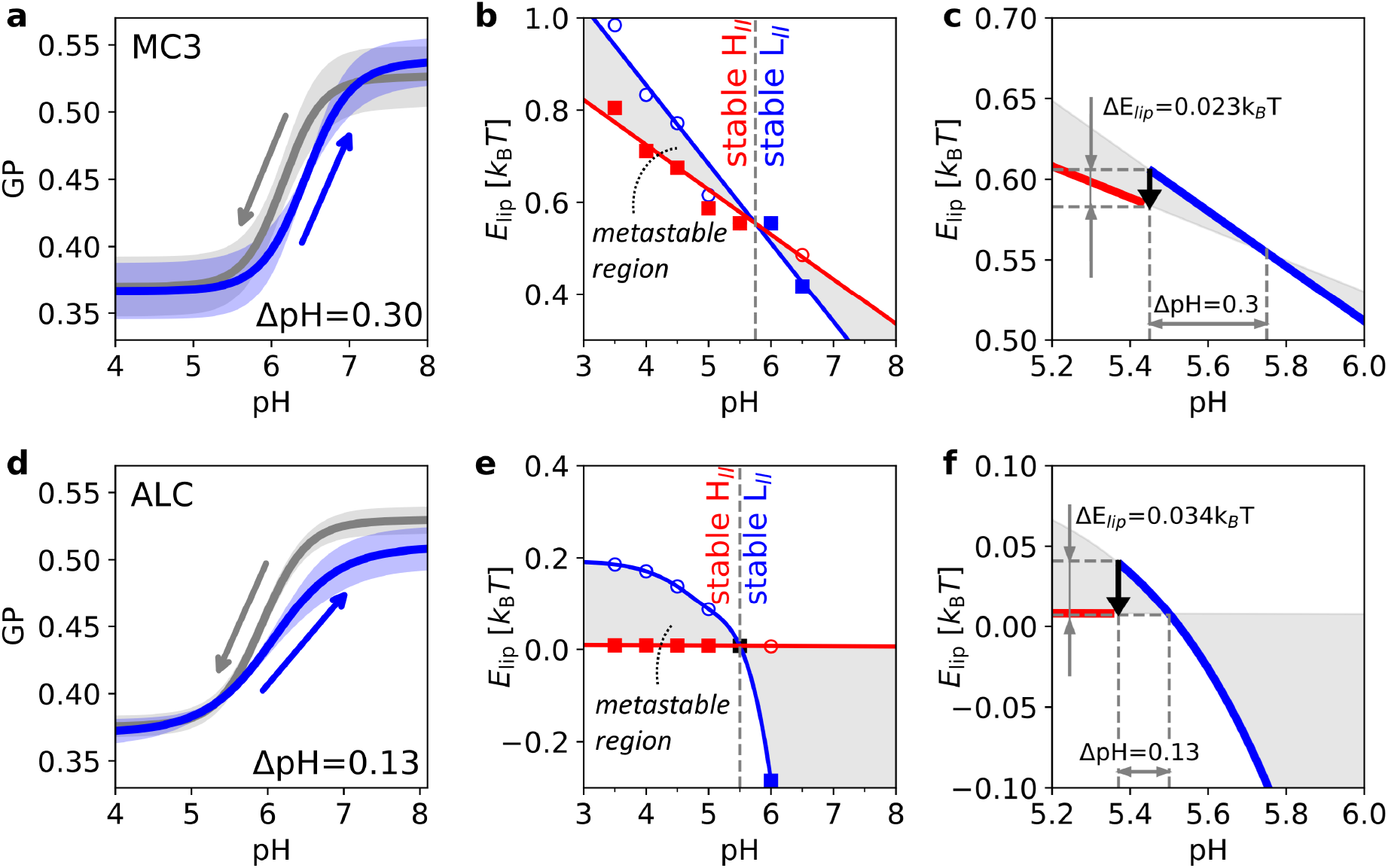
Hysteresis and metastable phase behavior in the L_*II*_–H_*II*_ transition **a**,**d.**Detection of hysteresis of the phase transition in MC3-containing (a) and ALC-containing (d) LNP measured with the Laurdan fluorescence. Arrows indicate the direction of pH titration. **b**,**e**. Energy per lipid of MC3/cholesterol (b) and ALC/cholesterol (e) system plotted as a function of pH. The continuum model determines the energy and next-neighbor distance as a function of spontaneous curvature. Using the experimentally established relationship between pH and next-neighbor distance, we map the experimental pH values onto the theoretical energy. The dashed line indicates the transition point and the colored area corresponds to a possible metastable region. Filled symbols indicate phases stable in that pH range and empty symbols indicate the unstable phases. Solid lines act as guide to the eye. **c**,**f**. Schematic representation of energy release when the transition happens from a metastable state. ΔpH values used for the calculation are taken from the GP measurement of hysteresis (figure 7 a,d).

## Conclusion

We establish a pH dependent continuum theory for mesophase transitions of ionizable lipids by combining SAXS, fluorescence spectroscopy, molecular dynamics simulations, and continuum theory. The close agreement in the pH- and temperature-dependent phase behavior of bulk systems and LNPs, as revealed by Laurdan fluorescence, supports the use of bulk mesophases as a model for investigating the thermodynamics of structural transitions within LNPs. We show, that a continuum description must account not only for curvature elasticity but also for the energetic cost of filling the interstitial voids to understand the transition between inverse micellar (L_*II*_) and inverse hexagonal (H_*II*_) phases, demonstrating that the energetic cost of filling the interstitial voids between curved lipid assemblies cannot be neglected. By integrating the theoretical model with experimental and simulation data, we obtain quantitative values for the elastic and structural properties of MC3 and ALC-0315 mesophases and their modulation by cholesterol. Despite their different molecular architectures, both ionizable lipids can be described within the same theoretical framework, linking molecular structure and composition to mesophase properties. The hysteresis of the L_*II*_–H_*II*_ transition upon pH cycling indicates that LNPs can become kinetically trapped in a metastable mesophase, providing evidence for an energy barrier between the two structural states. Together, these findings establish a quantitative link between the elastic properties of ionizable lipids and their pH-dependent mesophase behavior. This connection provides a physical basis for understanding how structural transitions and the associated energetic changes during endosomal acidification may contribute to membrane destabilization and LNP-mediated delivery.

## Acknowledgements

This work was supported by the Federal Ministry of Research, Technology and Space (BMFTR), Germany under Project 05K2026 - 2025-08197 LIPION within the framework of the Swedish–German research collaboration Röntgen-Ångström Cluster. We greatly acknowledge Deutsches Elektronen-Synchrotron (DESY) P12 BioSAXS EMBL and P62 SAXSMAT beamlines at PETRA III (Hamburg, Germany), and the beamline scientists Clement E. Blanchet and Andre Luiz Coelho Conceicao for the great support during the beamtimes.

## Methods

### Preparation of Bulk phases

Ionizable lipid bulk phases were prepared as described previously [philipp2023]. Briefly, ionizable lipids were dissolved in ethanol at a concentration of 48 mg/mL. Where applicable, cholesterol was added at an IL:cholesterol molar ratio of 3:1, and Laurdan was added at a concentration of 0.5 mM. The resulting solution was transferred into dialysis cups (Slide-A-Lyzer™ MINI, 3.5 kDa MWCO), with 215 *µ*L of lipid solution per cup, followed by the rapid addition of 80 *µ*L of citrate buffer (50 mM, pH 3). The samples were then dialyzed against citrate buffer (50 mM, pH 3) containing 25% ethanol for 48 h, followed by dialysis against PBS (1mM KH_2_PO_4_, 3mM Na_2_HPO_4_, 155 mM NaCl, pH 7.4) for 48 h and, finally, against citrate buffer adjusted to the desired pH. Afterwards, the bulk phases were transferred to glass capillaries for SAXS measurements and to eppendorf tubes for Laurdan fluorescent spectroscopy.

### LNP formulation

LNPs consisted of either DLin-MC3-DMA or ALC-315 as the ionizable lipid, together with DSPC, cholesterol, and DMPE-PEG2000, in a molar ratio of 50:10:38.5:1.5. Polyadenylic acid (PolyA), used as a surrogate for mRNA, was dissolved in sodium acetate buffer (25 mM, pH 4), while the individual lipid components were dissolved in ethanol (5 mM). LNPs were prepared by microfluidic mixing using a Herringbone mixer (Fluidic 1460, ChipShop) in combination with a Harvard Apparatus PHD 2000 syringe pump. The mixing was performed at a total flow rate (TFR) of 2 mL/min, a flow rate ratio (FRR) of 1:1 (aqueous:organic) for LNPs with DLin-MC3-DMA and TFR of 4 ml/min and FRR 3:1 for LNPs with ALC-315 and a N/P ratio of 6. Following a 20 min incubation at room temperature, LNPs were dialyzed against citrate buffer (50 mM citrate, 150 mM NaCl, pH 7.0) using Slide-A-Lyzer MAXI dialysis cups (3.5 kDa MWCO, Thermo Fisher Scientific) for at least 12 h at room temperature to remove residual ethanol. After LNP preparation, particle size distribution was determined by dynamic light scattering (DLS) using a Zetasizer Ultra Red (Malvern Panalytical), and encapsulation efficiency was assessed using the Quant-iT RiboGreen RNA assay (Invitrogen). The LNPs were subsequently adjusted to different pH conditions by buffer exchange using Slide-A-Lyzer MINI dialysis cups (3.5 kDa MWCO, Thermo Fisher Scientific) into citrate buffer (50 mM citrate, 150 mM NaCl; pH 4.0–7.5) for at least 3 h at room temperature. Lipid nanoparticles (LNPs) were incubated with 0.01 mg/ml Laurdan dissolved in DMSO at a volume ratio of 1:100 for 1 hour at 20 ^*°*^C and then transferred into Eppendorf tubes for GP measurement.

### SAXS measurements

Synchrotron small-angle X-ray scattering (SAXS) was carried out at the P12 EMBL BioSAXS and P62 SAXSMAT beamlines, PETRA III, DESY (Hamburg, Germany). The beamline instrumentation has been described previously.[37, 38] All the measurements were performed in borosilicate glass capillaries purchased from Hilgenberg. The bulk phases were always surrounded by excess of the correcponding buffer to avoid drying. The scattering data background was subtracted by measuring empty capillaries. The crystallographic space groups of the liquid crystalline phases and the nearest-neighbour distances (*d*_*NN*_) were determined from relative peak positions (see SI section 5).

### GP measurements of Bulk Phase and LNPs

The pH- and temperature-dependent structural states of bulk phases and LNPs were characterized by generalized polarization (GP) measurements using the solvatochromic fluorescent probe Laurdan (6-dodecanoyl-2-dimethylaminonaphthalene, Sigma-Aldrich) and a Lipid State Observer (LISO Pioneer, Lipospec GmbH), with GP values recorded over a temperature range of 10 to 55 ^*°*^C at a heating rate of 1 K/min and data collected at 1 K intervals.

Bulk phases with cholesterol and either DLin-MC3-DMA or ALC-315 were prepared with Laurdan at varying pH values as described above. The pre-stained sample phases were dispensed into an Eppendorf tube toward the detector, and the corresponding citrate buffer was added to the tube.

GP measurements of the bulk phase and LNPs were therefore performed as a function of temperature at different pH values. For the bulk phase, three samples from a single production batch were measured for each pH and ionizable lipid For LNPs, three independent productions were analyzed for each pH condition. Thus, three temperature scans were obtained for each pH value and averaged for further analysis.

To determine the GP as a function of pH, the corresponding GP values were extracted from the temperature scans at defined temperatures. At each temperature, the pH-dependent GP values were fitted with a sigmoidal function in OriginLab, with the inflection point defining the phase transition pH* and the transition width representing its uncertainty (Figure S13).

To measure the hysteresis of the phase transition in LNPs, four LNP production batches were prepared for each ionizable lipid, as described above. The pH was reduced by stepwise adding HCl (0.1 and 1 M) and monitored using a pH meter (Lab 845, SI Analytics) equipped with a pH electrode (Biotrode 3 mm, METTLER TOLEDO). After each pH adjustment, a sufficient equilibration period (*∼* 1-2 min) was allowed before GP values were recorded in triplicate using the Lipid State Observer. Once a pH of 4 was reached, the process was repeated by adding NaOH (0.1 and 1 M) in steps until the initial pH was reached. Each set of measurements was fitted to a sigmoid function in OriginLab, and the mean and standard deviation of the four different LNP productions were plotted (Figure S12). The pH* value of the phase transition was then determined based on the inflection point.

### MD Simulations

To obtain atomistic insights into the structural and bending properties of various H_*II*_ phases, all-atom MD simulations were performed using the GROMACS software package (version 2024.4) [39]. Packmol [40] was employed to generate the starting configurations for the simulations. A total of 4 distinct H_*II*_ phases were modeled: (i) Pure MC3, (ii) MC3/cholesterol (3:1) molar ratio, (iii) pure ALC and (iv) ALC/cholesterol (3:1 molar ratio). A triclinic box was employed to replicate the symmetry of the H_*II*_ phases with the water channel oriented parallel to the Z axis. All the ionizable lipids were assumed to be fully protonated. To mimic physiological conditions, the system was solvated with a salt concentration of 0.15 M and additional counter ions were added to neutralize the charged nature of the ionizable lipids. Water content was iteratively adjusted to reproduce the experimentally observed lattice spacings from small-angle X-ray scattering (SAXS) data. The composition of each H_*II*_ phase is summarized in Table 2. The force field parameters for MC3 were adopted from our previous study [41], ALC parameters were obtained from Paloncýová et al. [42] and the parameters for cholesterol were derived from the Lipid17 AMBER force field [43]. The Mamatkulov-Schwierz ion parameters [44] were used along with TIP3P water model [45] for MC3-based H_*II*_ phases and OPC water model [46] for ALC-based H_*II*_ phases.

**Table 2:** Composition of simulated H_*II*_ phases. The lattice spacing (*d*_*NN*_) is matched to the experimental SAXS data. *n*_*w*_ represents the water to lipid ratio required to maintain the target spacing.

| System | Ionizable lipid | Cholesterol | Water | Na<sup>+</sup> | Cl<sup>-</sup> | $n_w$ | $d_{NN}$ (nm) |
| --- | --- | --- | --- | --- | --- | --- | --- |
| MC3 w/o chol | 200 | 0 | 3600 | 10 | 210 | 18 | 6.42 |
| MC3 with chol | 150 | 50 | 2400 | 7 | 157 | 12 | 6.00 |
| ALC w/o chol | 200 | 0 | 2400 | 7 | 207 | 14 | 4.41 |
| ALC with chol | 150 | 50 | 1600 | 4 | 154 | 8 | 4.37 |

The H_*II*_ phases were first subjected to steepest descent energy minimization, followed by a short equilibration phase to assign velocities and stabilize the system. The system was first equilibrated for 2 ns under semi-isotropic pressure coupling using the Berendsen barostat with a time constant of 1 ps to stabilize box dimensions, followed by a 200 ns equilibration using the Parrinello–Rahman barostat with a time constant of 5 ps. Temperature was maintained at 298 K using a Nosé-Hoover thermostat with a time constant of 1ps, with independent coupling of the lipid membrane and the solvent including ions. Finally, production simulations of 200 ns were carried out under NVT conditions using a 2 fs time step. Short-range electrostatic and van der Waals interactions were evaluated with a cutoff of 1.2 nm, while long-range electrostatic interactions were treated using PME with a fourier grid spacing of 0.12 nm.and pme interpolation order of 4.

All hydrogen-containing bonds were constrained using the LINCS algorithm and non-bonded interactions were computed on-the-fly with Verlet neighbor lists with a cut-off of 1.2 nm updated every 20 steps. Analysis of the simulation trajectory was done with MDAnalysis [47] scripts and visualization of the trajectory and generation of simulation snapshots were done with VMD [48]. A detailed description of the bending rigidity calculation is provided in the Supporting Information (Section 5.1).

