## Supplementary Information for "Thermodynamic Model and Experiments Indicate Kinetically Trapped States in Ionisable Lipid Nanoparticles"

#### Contents

**1 Data Fitting** **2**
1.1 Fit of MC3 and ALC-0315 data without cholesterol . . . . . . . . . . . . . 3

**2 Scaling of the void energy term** **4**

**3 Estimating the water content** **4**

**4 Transition energy from hysteresis in the $L_{II}$-$H_{II}$ transition** **5**

**5 SAXS data** **8**

### 1 Data Fitting

SAXS measurements yield the nearest-neighbor distances of the micellar and hexagonal phases. Combined with the offset $\iota$ between the nearest-neighbor distance and the pivotal plane position obtained from MD simulations, the curvature radius is determined as ($R_C = d_{NN}^{(H_{II})}/2-\iota$) for the hexagonal phase and ($R_S = d_{NN}^{(L_{II})}/2-\iota$), respectively. MD simulations additionally provide the bending rigidity $\kappa$. The remaining fitting parameters are the Gaussian bending rigidity, $\kappa_G$, the void parameters, $\varepsilon^{(H_{II})}$, $\varepsilon^{(L_{II})}$, and the spontaneous curvature $R_0$. Whereas $R_0$ depends on protonation state and thus pH, $\kappa_G$, $\varepsilon^{(H_{II})}$, and $\varepsilon^{(L_{II})}$ are assumed to be independent of pH.

Furthermore, $\kappa$ is known from MD simulations. For a fixed pH value, $R_C$ and $R_S$, respectively adjust to minimise the energy. For zero pressure, the condition $\frac{dE_{tot}^{H_{II}}}{dR_C} = 0$ allows to express $\varepsilon^{(H_{II})}$ in dependence of $R_C$, $R_0$ and $\kappa_G/\kappa$ :

$$\frac{\varepsilon^{(H_{II})}}{\kappa} = \frac{1}{R_C^3}\left(\frac{1}{2R_C} - \frac{1}{R_0}\right) \tag{M1}$$

Similarly, $\frac{dE_{tot}^{L_{II}}}{dR_S} = 0$ leads to

$$\frac{\varepsilon^{(L_{II})}}{\kappa} = \frac{2}{R_C^3}\left(\frac{1}{R_C} - \frac{1}{R_0}\right) + \frac{\kappa_G}{\kappa}\frac{1}{R_S^4} \tag{M2}$$

At the mesophase transition for which we denote the curvature radii as $R_C^*$ and $R_S^*$ the energy of both mesophases has to be equal, which leads to an expression for the spontaneous curvature radius $R_0^*$ at the transition, that is expressed in dependence of only $R_C^*$ , $R_S^*$ and $\kappa_G/\kappa$:

$$R_0^* = \frac{1/R_C^* - 6/R_S^*}{1/R_C^{*2} - 4/R_S^{*2} - 2\kappa_G/(\kappa R_S^{*2})} \tag{M3}$$

Hence $R_0^*$ can be expressed in terms of quantities that are determined from experiment and MD simulations and the so far unknown $\kappa_G$.

Since Equs. M1 and M2 hold for all pH values they also hold at the transition point. Hence, $\varepsilon^{(H_{II})}$, $\varepsilon^{(L_{II})}$ are also be expressed in terms of known quantities and $\kappa_G/\kappa$.

We now turn to non-zero osmotic pressure. Considering a pH-value close the transition point in the $H_{II}$ phase and write the pressure as

$$p = 4\kappa\left[\left(\frac{1}{2R_C} - \frac{1}{R_0^*(\kappa_G/\kappa)}\right)\frac{1}{R_C^2} - \frac{\varepsilon^{(H_{II})}(\kappa_G/\kappa)}{\kappa}R_C\right]. \tag{M4}$$

We note, that the spontaneous curvature is considered to be determined primarily by the protonation state of the ionizable lipids and is thus pressure independent. In contrast, the curvature radii $R_C$ and $R_S$ adjust to the applied pressure. As in the zero-pressure case, $R_C$ and $R_S$ are determined combing knowledge from SAXS measurements and MD simulation.

#### 1.1 Fit of MC3 and ALC-0315 data without cholesterol

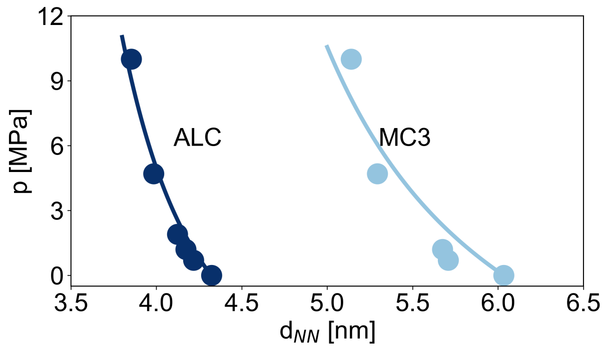

Figure S1: Next-neighbour distance as a function of osmotic pressure for MC3 mesophases at pH 5.5, and ALC-0315 mesophases at pH 5.0, in both cases w/o cholesterol. Solid lines show the best fit with the theoretical model (eq. 7)

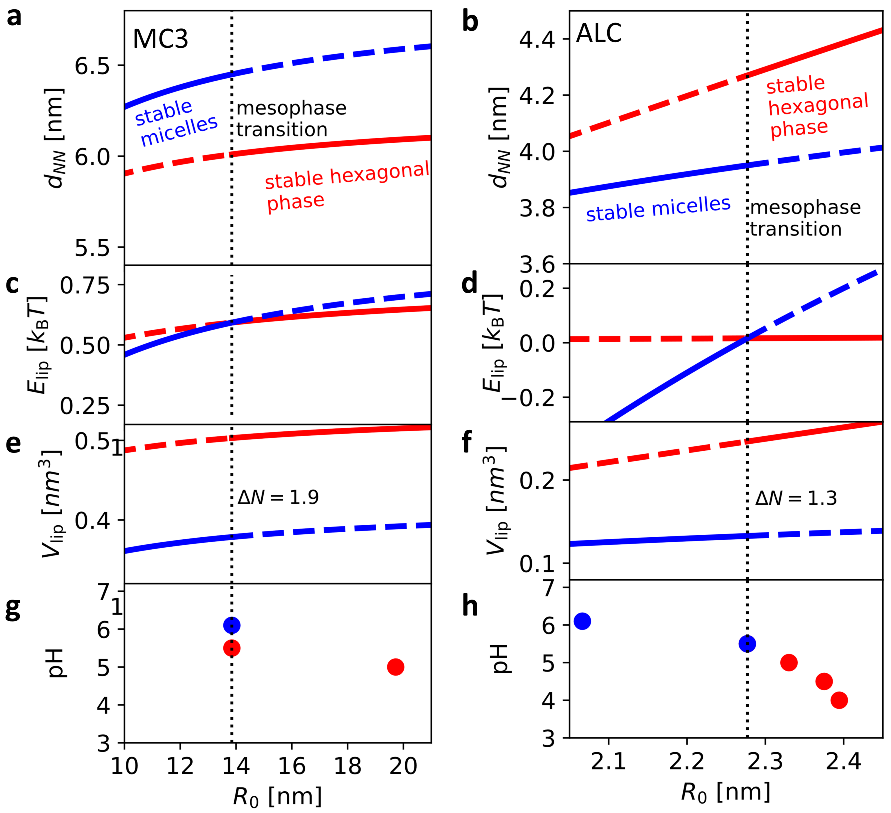

Figure S2: Numerical calculation of lipid free energy and volume per lipid as a function of spontaneous curvature and pH. The curves are obtained using the fitted parameters (Table 1) for the phase transition for MC3 (left column) and ALC (right column) systems without cholesterol. **a,b.** Fitted dependence of next-neighbour distance ($d_{NN}$) on the spontaneous curvature radius ($R_0$), **c,d.** Calculated energy per lipid ($E_{lip}$) as a function of $R_0$, **e,f.** Calculated volume per lipid ($V_{lip}$) as a function of $R_0$, **g,h.** $R_0$ as a function of pH obtained using experimental $d_{NN}$(pH) data

#### 2 Scaling of the void energy term

In this section, we justify the scaling form used for the void-filling energy. We show that both lipid stretching and cholesterol redistribution give rise to the same $E_{void} \propto A_{tot}R^2$ dependence. In both cases, the number of molecules participating in void filling is proportional to the total interfacial area $A_{tot}$, since the interface consists of a fixed areal density of lipids (and associated cholesterol). The energetic cost can therefore be written as an energy per molecule multiplied by $A_{tot}$.

For lipid stretching, the mismatch that must be accommodated by each lipid increases with the characteristic radius $R$. The required tail deformation is therefore proportional to $R$. Assuming a harmonic elastic response, the energy per lipid scales as $R^2$, and thus $E_{void}^{stretch} \sim A_{tot}R^2$.

For cholesterol redistribution, the amount of cholesterol required to fill the void scales with the void volume, which in turn scales as $A_{tot}R$. Since the number of lipids and associated cholesterol sites at the interface scales with $A_{tot}$, the required change in local composition per lipid scales as $\delta c \propto R$. Assuming a harmonic free-energy cost for deviations from the preferred composition, the mixing free energy scales as $f \propto (\delta c)^2 \propto R^2$. Multiplying by the number of interfacial molecules ($\propto A_{tot}$) gives $E_{void}^{chol} \propto A_{tot}R^2$, yielding the same $R^2 A_{tot}$ scaling as for lipid stretching. Thus, both mechanisms give scaling behavior

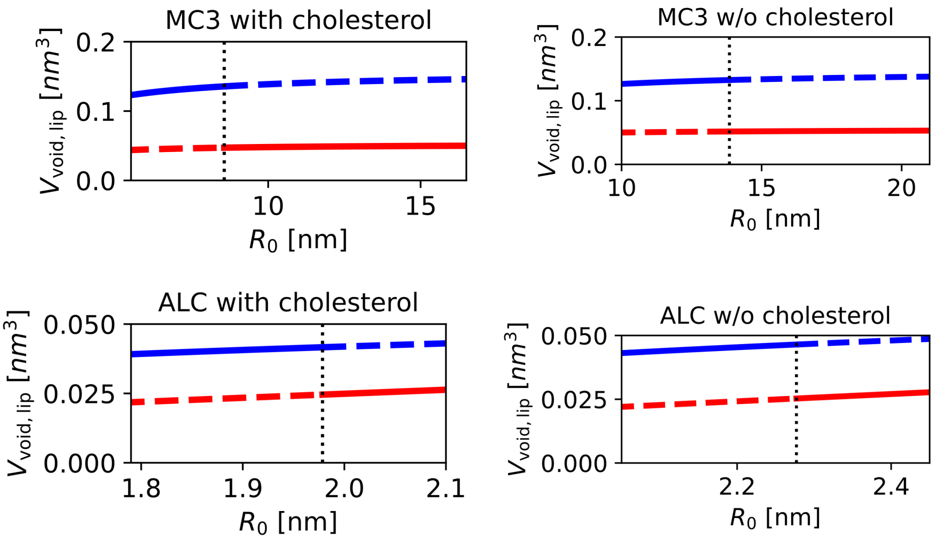

Figure S3: Calculation of the void volume per lipid as a function of spontaneous curvature radius for MC3 and ALC systems with and without cholesterol.

#### 3 Estimating the water content

In the following, it is derived how the volume and water content change as the lipid phase transitions from the $L_{II}$ to the $H_{II}$ phase.

In the $L_{II}$ phase the area per micelle is given by

$$\begin{aligned} A_S &= 4\pi R_S^2 \\ &= N_{lipid} a_{lip} \, , \end{aligned} \tag{1}$$

with $R_\mathrm{S}$ the curvature radius, $N_\mathrm{lipid}$ the number of lipids per micelle and $a_\mathrm{lip}$ the area per lipid. The volume of a micelle is then obtained as

$$\begin{aligned} V_\mathrm{S} &= \frac{4}{3}\pi R_\mathrm{S}^3 \\ &= N_\mathrm{lipid}\frac{a_\mathrm{lip}R_\mathrm{S}}{3}\,. \end{aligned} \tag{2}$$

We now consider a section of cylinder in the $\mathrm{H_{II}}$ phase with the same number of lipids as the micelle. The area then reads

$$\begin{aligned} A_\mathrm{C} &= 2\pi R_\mathrm{C} L \\ &= N_\mathrm{lipid} a_\mathrm{lip}\,, \end{aligned} \tag{3}$$

with $L$ the length of the cylinder. Similarly, the volume is obtained as

$$\begin{aligned} V_\mathrm{C} &= 2\pi R_\mathrm{C}^2 L \\ &= N_\mathrm{lipid}\frac{a_\mathrm{lip}R_\mathrm{C}}{2}\,. \end{aligned} \tag{4}$$

Thus, the volume per lipid $V_\mathrm{lip}$ reads

$$V_\mathrm{lip} = \begin{cases} \frac{a_\mathrm{lip}R_\mathrm{S}}{3}\,; & \mathrm{L_{II}}\ \text{phase} \\ \frac{a_\mathrm{lip}R_\mathrm{C}}{2}\,; & \mathrm{H_{II}}\ \text{phase}\,. \end{cases} \tag{5}$$

To estimate the change in water content, we assume that the relative change in water content is the same as the relative change in volume:

$$\frac{\Delta N_\mathrm{wat}}{N_\mathrm{wat}} = \frac{V_\mathrm{C} - V_\mathrm{S}}{V_\mathrm{C}}\,, \tag{6}$$

with $N_\mathrm{wat}$ the number of water molecules per lipid in the $\mathrm{H_{II}}$ phase and $\Delta N_\mathrm{wat}$ the change of number of water molecules per lipid. This expression is equivalent to

$$\Delta N_\mathrm{wat} = \left(1 - \frac{2}{3}\frac{R_\mathrm{S}}{R_\mathrm{C}}\right) N_\mathrm{wat}\,. \tag{7}$$

Based on a previous work, we assume a number of water molecules per lipid of $N_\mathrm{wat} = 12$ for MC3 and $N_\mathrm{wat} = 8$ for ALC [1].

### 4 Transition energy from hysteresis in the $\mathrm{L_{II}}$-$\mathrm{H_{II}}$ transition

The change in GP with increasing/decreasing pH is used to determine the energy difference $\Delta E_\mathrm{trans}$ to initial the mesophase transition. We first consider the case of decreasing pH and note that an analog derivation can be done for increaing pH. The pH-dependent GP is describes as

$$GP(pH) = GP_\mathrm{H_{II}} + (GP_\mathrm{L_{II}} - GP_\mathrm{H_{II}})p_\mathrm{L_{II}}(pH)\,, \tag{8}$$

with $p_\mathrm{L_{II}}(pH)$ is the probability of being in the $\mathrm{L_{II}}$ phase and $GP_\mathrm{L_{II}}$, $GP_\mathrm{H_{II}}$ the GP signal in the pure $\mathrm{L_{II}}$, $\mathrm{H_{II}}$ phase. In a three-state system, the probability $p_\mathrm{L_{II}}$ is related to the energy $E_{\mathrm{L_{II}}}$ in the pure $\mathrm{L_{II}}$ phase, the energy of the intermediate structure $E_{\mathrm{inter}}$ and the energy of the pure $\mathrm{H_{II}}$ phase as

$$p_{\mathrm{L_{II}}} = \frac{\exp[-E_{\mathrm{L_{II}}}/k_{\mathrm{B}}T]}{\exp[-E_{\mathrm{L_{II}}}/k_{\mathrm{B}}T] + \exp[-E_{\mathrm{inter}}/k_{\mathrm{B}}T] + \exp[-E_{\mathrm{H_{II}}}/k_{\mathrm{B}}T]}\,, \tag{9}$$

with $k_{\mathrm{B}}T$ the thermal energy. Similarly, we write the probability to be in an intermediate structure $p_{\mathrm{inter}}$ or the pure $\mathrm{H_{II}}$ phase $p_{\mathrm{H_{II}}}$ as

$$p_{\mathrm{inter}} = \frac{\exp[-E_{\mathrm{inter}}/k_{\mathrm{B}}T]}{\exp[-E_{\mathrm{L_{II}}}/k_{\mathrm{B}}T] + \exp[-E_{\mathrm{inter}}/k_{\mathrm{B}}T] + \exp[-E_{\mathrm{H_{II}}}/k_{\mathrm{B}}T]} \tag{10}$$

$$p_{\mathrm{H_{II}}} = \frac{\exp[-E_{\mathrm{H_{II}}}/k_{\mathrm{B}}T]}{\exp[-E_{\mathrm{L_{II}}}/k_{\mathrm{B}}T] + \exp[-E_{\mathrm{inter}}/k_{\mathrm{B}}T] + \exp[-E_{\mathrm{H_{II}}}/k_{\mathrm{B}}T]}. \tag{11}$$

Since all probabilities have to add up to 1, we write

$$\begin{aligned} 1 - p_{\mathrm{L_{II}}} &= p_{\mathrm{inter}} + p_{\mathrm{H_{II}}} \qquad (12)\\ &= \frac{\exp[-E_{\mathrm{inter}}/k_{\mathrm{B}}T] + \exp[-E_{\mathrm{H_{II}}}/k_{\mathrm{B}}T]}{\exp[-E_{\mathrm{L_{II}}}/k_{\mathrm{B}}T] + \exp[-E_{\mathrm{inter}}/k_{\mathrm{B}}T] + \exp[-E_{\mathrm{H_{II}}}/k_{\mathrm{B}}T]}\,. \end{aligned}$$

Hence, we find

$$\frac{p_{\mathrm{L_{II}}}}{1 - p_{\mathrm{L_{II}}}} = e^{-\Delta E_{\mathrm{inter}}/k_{\mathrm{B}}T} + e^{-(E_{\mathrm{L_{II}}} - E_{\mathrm{H_{II}}})/k_{\mathrm{B}}T}\,, \tag{13}$$

with $\Delta E_{\mathrm{int}} = E_{\mathrm{L_{II}}} - E_{\mathrm{inter}}$. For $E_{\mathrm{L_{II}}} - E_{\mathrm{H_{II}}} \gg \Delta E_{\mathrm{int}}$ this expression simlifies to

$$\frac{p_{\mathrm{L_{II}}}}{1 - p_{\mathrm{L_{II}}}} \approx e^{-\Delta E_{\mathrm{inter}}/k_{\mathrm{B}}T}\,. \tag{14}$$

Thus, we are able to estimate $\Delta E_{\mathrm{int}}$ from the experimental measurements using

$$\Delta E_{\mathrm{int}}(pH) = k_{\mathrm{B}}T \ln\left(\frac{GP_{\mathrm{L_{II}}} - GP(pH)}{GP(pH) - GP_{\mathrm{H_{II}}}}\right)\,. \tag{15}$$

As mentioned above, the analogous derivation can also be performed for increasing pH. Fig. S4a) shows the energy difference over the entire experimentally investigated pH range. Subfigure b) shows the region around the transition pH value together with the slope at $\Delta E_{\mathrm{int}} = 0$. We consider the two branches of the hysteresis curve as limiting cases that enclose the equilibrium transition curve. In all cases, the energy changes by approximately $|d\Delta E_{\mathrm{int}}/d\mathrm{pH}| \sim (2-4)\mathrm{k_B T}$ per unit pH around the transition midpoint. We estimate the transition energy that accounts for the system remaining in an energetically unfavorable phase in a small pH-regime around the transiton pH as $\Delta E_{\mathrm{trans}} = |d\Delta E_{\mathrm{int}}/d\mathrm{pH}|\Delta\mathrm{pH} \sim (0.5-1)\,\mathrm{k_B T}$.

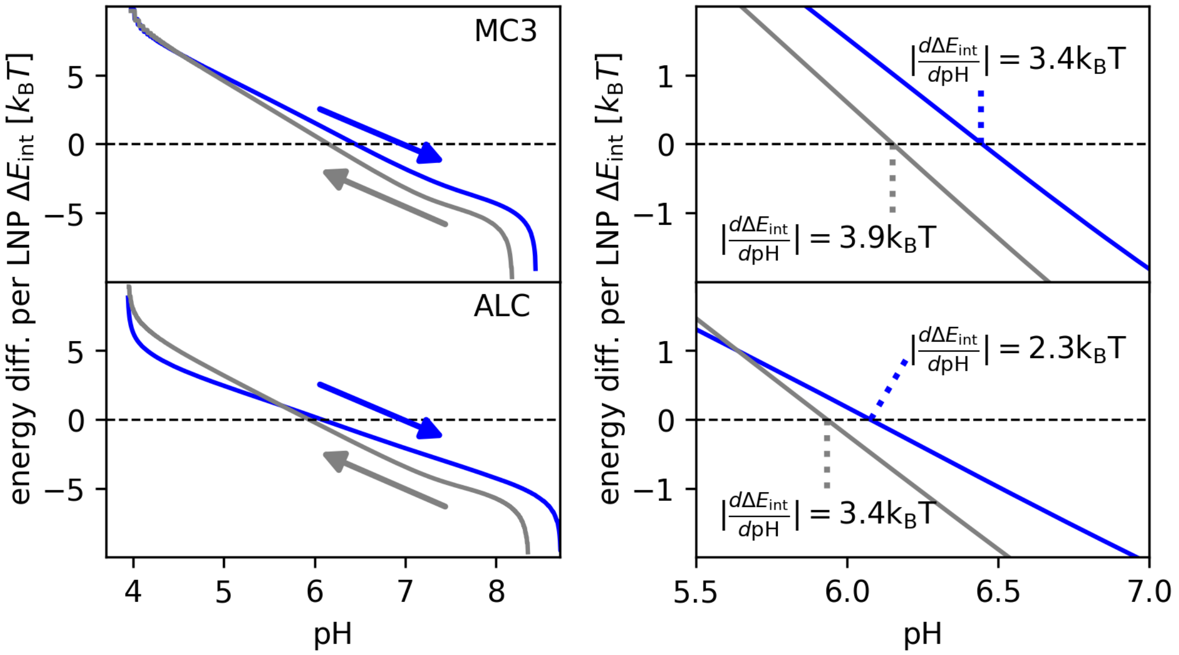

Figure S4: Energy difference $\Delta E_{\text{int}}$ to the transition state from fluorescence measurements: a) $\Delta E_{\text{int}}$ from Eq. 15 for the entire experimentally investigated pH range for both increasing (blue) and decreasing (grey) pH. b) $\Delta E_{\text{int}}$ in the vicinity of the transition pH.

### 5 SAXS data

#### 5.1 Bulk phases with and w/o cholesterol

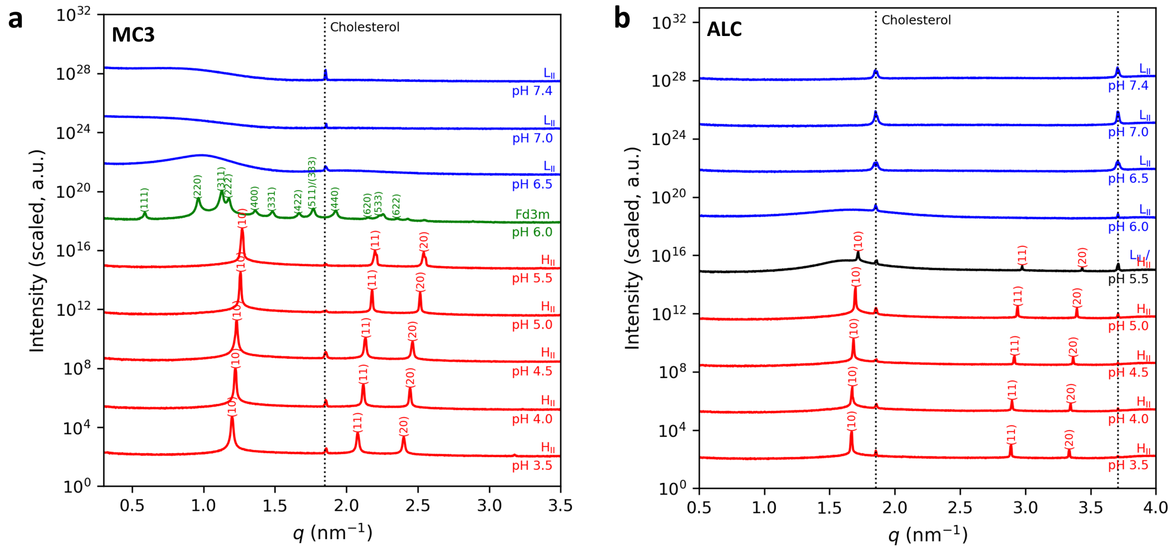

Figure S5: SAXS data of MC3/cholesterol (a) and ALC-0315/cholesterol (b) mesophases at different pH. All the crystalline peaks are indexed. Fd3m phase is formed from inverted micelles; the LII phase corresponds to the inverted micellar phase in liquid order. The cholesterol peak corresponds to cholesterol crystals formed from the excess of cholesterol in the system.

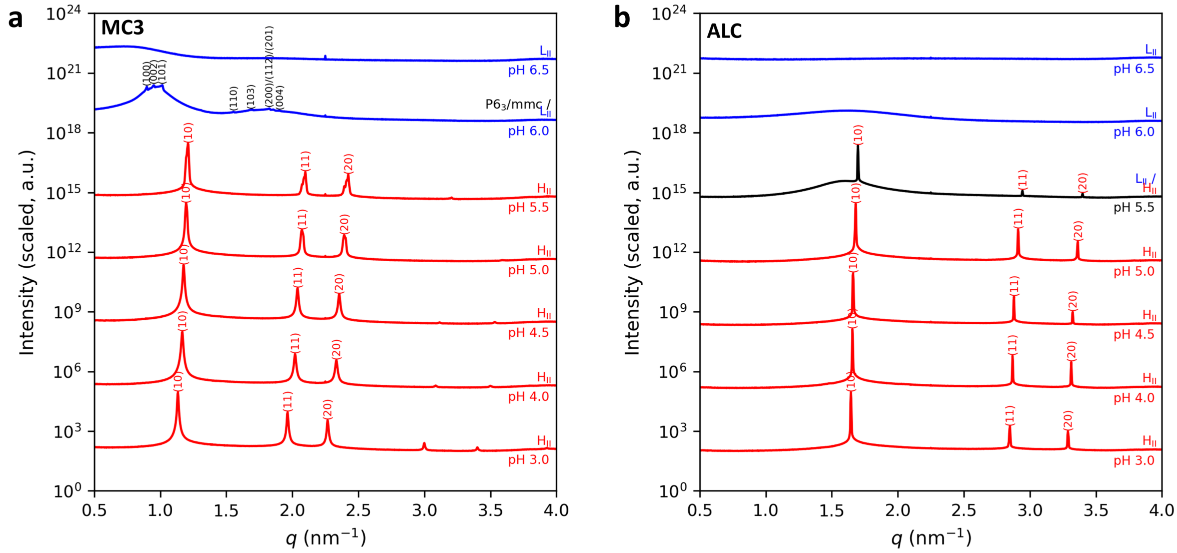

Figure S6: SAXS data of MC3 (a) and ALC-0315 (b) mesophases without cholesterol at different pH. All the crystalline peaks are indexed. $P6_3/mmc$ phase is formed from inverted micelles; the LII phase corresponds to the inverted micellar phase in liquid order. The cholesterol peak corresponds to cholesterol crystals formed from the excess of cholesterol in the system.

#### 5.2 Temperature data

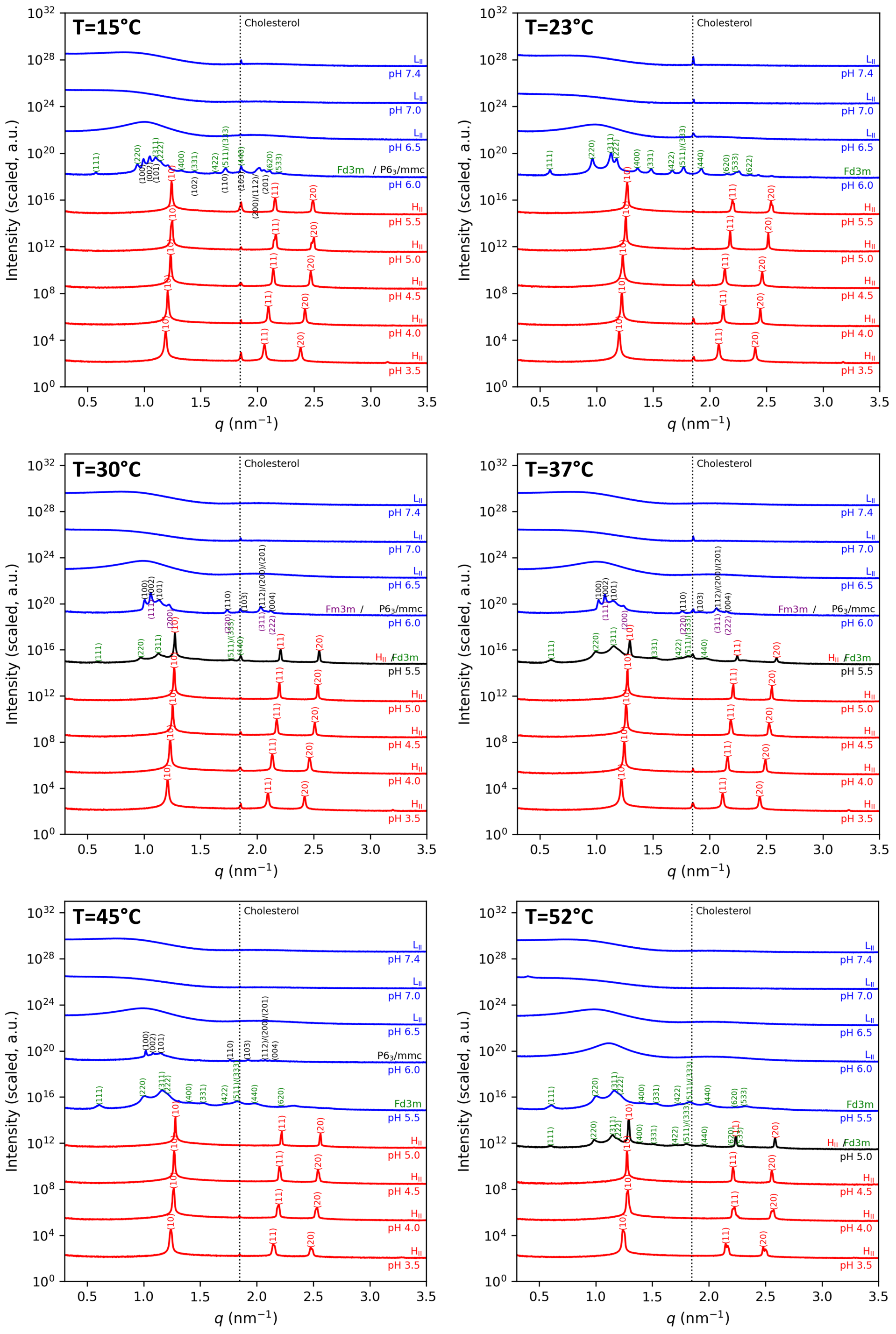

Figure S7: SAXS data of MC3/cholesterol mesophases at different pH and temperatures. All the crystalline peaks are indexed. Fd3m, Fm3m and $P6_3/mmc$ phases are formed from inverted micelles; the LII phase corresponds to the inverted micellar phase in liquid order. Cholesterol peak corresponds to cholesterol crystals formed from the excess of cholesterol in the system.

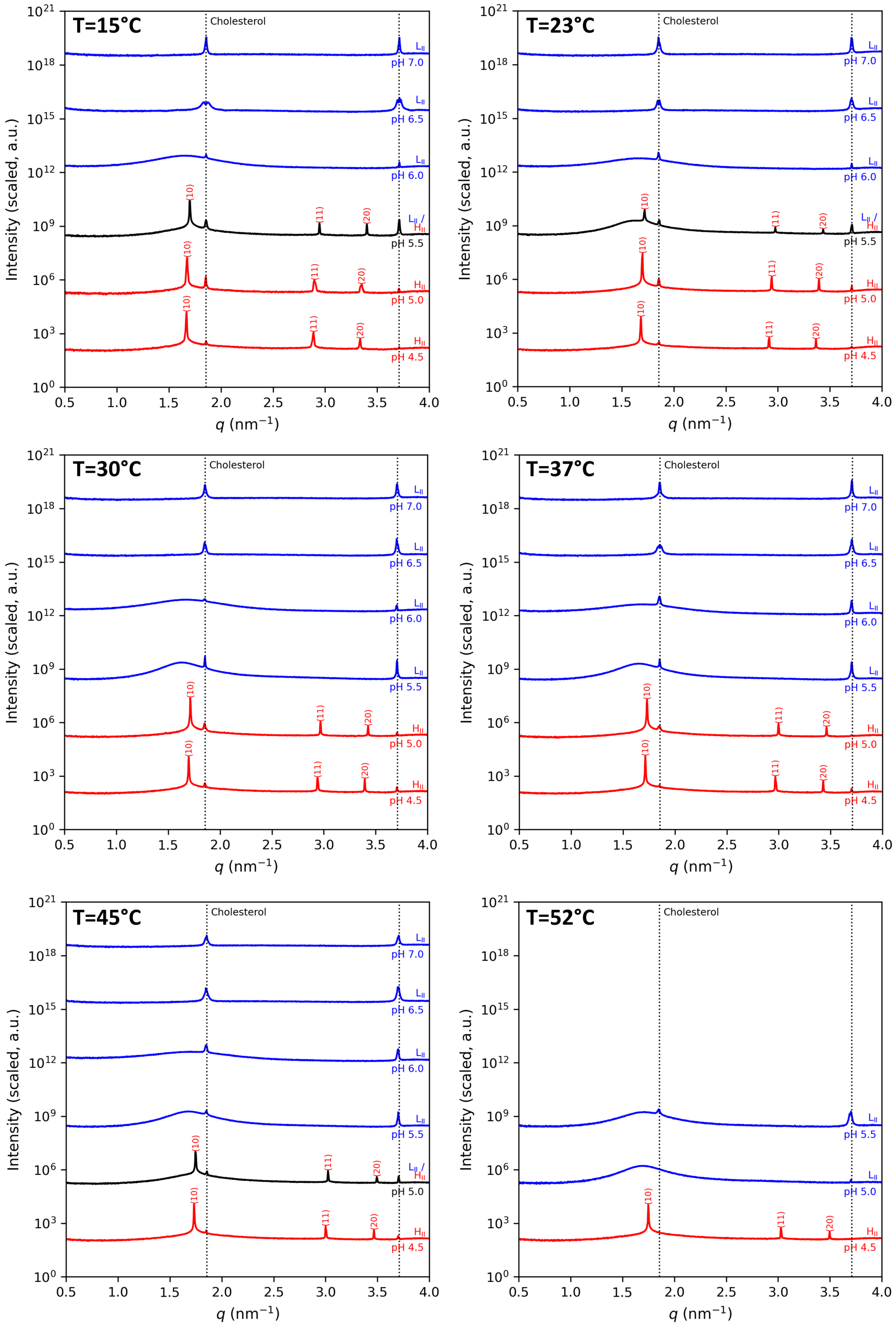

Figure S8: SAXS data of ALC/cholesterol mesophases at different pH and temperatures. The LII phase corresponds to the inverted micellar phase in liquid order. The cholesterol peak corresponds to cholesterol crystals formed from the excess of cholesterol in the system.

### 6 MD Simulations

#### 6.1 Calculation of bending rigidity

The bending rigidity ($\kappa$) of inverse hexagonal ($H_{II}$) lipid phases was determined from the distribution of lipid splay (S) computed from molecular dynamics (MD) simulation trajectories by adapting the methodology by Khelashvili and coworkers [2, 3]. The procedure comprises of (i) defining the geometric descriptors for the lipids, (ii) computing pairwise splay $S$ for neighboring lipids, and (iii) extracting $\kappa$ from the splay distribution.

The central water column of the $H_{II}$ phase oriented along the $Z$ axis provides a reference frame for defining lipid geometric descriptors essential to quantifying the bending rigidity of $H_{II}$ phases. For each simulation frame, the box length along the $Z$ direction ($L_Z$) was divided into 50 equally spaced bins. The local center of mass coordinates of the water molecules within each bin was computed and the geometric descriptors of the lipids within each bin were then defined relative to the local water center of mass.

**Lipid geometric descriptors** For each lipid in the system, three geometric descriptors were defined to calculate the splay distribution (Fig. S8):

1. **Pivotal plane**: The pivotal plane is obtained by systematically evaluating the area per lipid $A_L$ as a function of the average radial atomic positions of the lipid monolayer. This is repeated across multiple systems with varying water content, enabling a direct assessment of the response of $A_L$ to changes in the membrane curvature. The radial positions where $A_L$ remains invariant across multiple membrane curvatures is chosen as the pivotal plane. The area per lipid $A_L$ at the pivotal plane was calculated from the pivotal plane distance $R_p$ and the simulation box height $L_z$:

$$A_L = \frac{2\pi R_p L_z}{N_{\text{lipids}}} \tag{16}$$

   where $N_{\text{lipids}}$ is the total number of lipids in the system.

2. **Lipid director vector** $\hat{n}$: Unit vector pointing from the tail group to the head group, defining the local orientation of the lipid.

3. **Surface normal vector** $\hat{N}$: Defined as the normalized vector from the position of the pivotal plane to the local water center of mass.

**Pairwise splay calculation** For each frame, the pairwise splay between neighboring lipid pairs separated by a cut-off distance of 10 $\AA$ was computed based on the procedure in [2, 3]. The separation vector between neighboring lipids $i$, $j$ was projected onto the local tangent plane of lipid i to remove the component along the surface normal.

$$\mathbf{r}_{ij}^{\parallel} = \mathbf{r}_{ij} - (\mathbf{r}_{ij} \cdot \hat{\mathbf{N}}_i)\,\hat{\mathbf{N}}_i \tag{17}$$

The projected separation was normalized to obtain the unit tangent vector: $\hat{\mathbf{r}}_{ij} = \mathbf{r}_{ij}^{\parallel}/\|\mathbf{r}_{ij}^{\parallel}\|$. The splay $S_{ij}$ for the pair was computed as:

$$S_{ij} = \frac{(\hat{\mathbf{n}}_j - \hat{\mathbf{n}}_i) \cdot \hat{\mathbf{r}}_{ij}}{\|\mathbf{r}_{ij}\|} - \frac{(\hat{\mathbf{N}}_j - \hat{\mathbf{N}}_i) \cdot \hat{\mathbf{r}}_{ij}}{\|\mathbf{r}_{ij}\|} \tag{18}$$

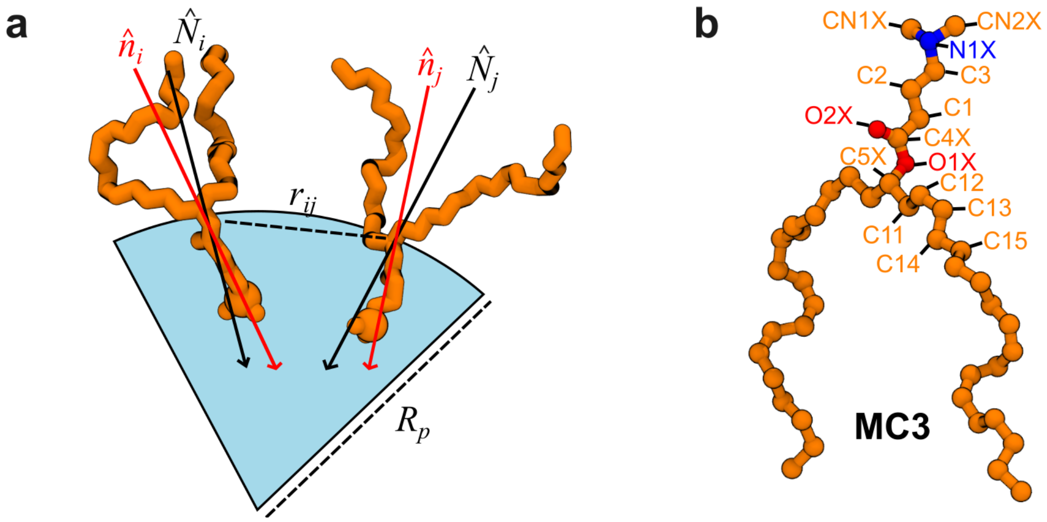

Figure S9: (a) Geometric descriptors used to compute the splay between neighboring lipids in the $H_{II}$ phase. $\hat{n}_i$ and $\hat{n}_j$ represent the lipid director vectors, $\hat{N}_i$ and $\hat{N}_j$ represent the local surface normal vectors at the pivotal plane, $r_{ij}$ is the distance between the pivotal atoms of both lipids and $R_p$ is the radial distance from the center of the water channel to the pivotal plane. (b) Name of the individual atoms of MC3 used in the calculation of the position of the pivotal plane in Fig.4 in the main text. The carbon, nitrogen and oxygen atoms are colored orange, blue and red respectively.

where $r_{ij}$ is the distance between the pivotal atoms of both the lipids.

In the case of $H_{II}$ phases with multiple lipid compositions, splay values were accumulated and binned by lipid pair type (e.g., MC3–MC3, Chol–Chol, MC3–Chol) to enable the computation of pair-specific bending rigidities.

**Extracting bending rigidity from splay distribution** The energetic cost of deforming a membrane from its preferred curvature is described by Helfrich elastic theory, with the local deformation energy per unit area given by:

$$f = \frac{1}{2}\kappa S^2 \tag{19}$$

where $S$ denotes the local splay and $\kappa$ is the bending rigidity. The distribution of splay in a small patch of the membrane follows a Boltzmann distribution:

$$P(S) \propto \exp\left[-\frac{\kappa A_L}{2k_BT}S^2\right]. \tag{20}$$

The potential of mean force (PMF) associated with the splay was obtained from:

$$\frac{\mathrm{PMF}(S)}{k_BT} = -\ln P(S) = \frac{\kappa A_L}{2k_BT}S^2 + \mathrm{const} \tag{21}$$

Within the harmonic approximation, the PMF is parabolic near its minimum:

$$\frac{\mathrm{PMF}(S)}{k_BT} = b\,(S - S_0)^2 + \epsilon \tag{22}$$

where $b$ is the curvature of the PMF, $S_0$ its minimum, and $\epsilon$ an offset. By fitting this quadratic form to the PMF data within the range ($-0.05 \leq S \leq 0.05$ Å$^{-1}$), the bending rigidity in units of $k_BT$ was extracted:

$$\frac{\kappa}{k_BT} = \frac{2b}{A_L} \tag{23}$$

with uncertainty $\delta\kappa = 2\,\delta b/A_L$.

**Multi-Component Membranes** For membranes containing multiple lipid species (e.g., MC3/cholesterol or ALC/cholesterol mixtures), bending rigidity was first computed independently for each lipid pair type ($\kappa_{\alpha\beta}$ for pair $\alpha$–$\beta$). The effective bending rigidity $\kappa_{\text{eff}}$ for the multi-component membrane was then obtained as a pair-count-weighted average following the procedure in [2, 3]:

$$\kappa_{\text{eff}} = \left[\frac{1}{\sum_{\alpha\leq\beta} N_{\alpha\beta}} \sum_{\alpha\leq\beta} \frac{N_{\alpha\beta}}{\kappa_{\alpha\beta}}\right]^{-1} \tag{24}$$

where $N_{\alpha\beta}$ is the total number of pair interactions of type $\alpha$–$\beta$ across all analysed frames.

#### 6.2 $\kappa$ for ALC-Cholesterol $H_{II}$ phase

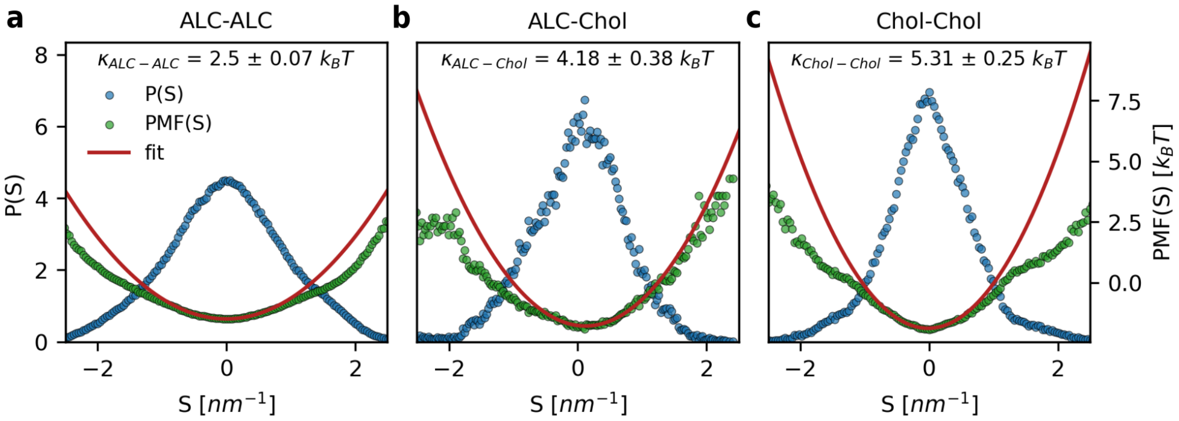

Figure S10: Bending rigidity $\kappa$ obtained from ALC-Cholesterol $H_{II}$ phases for (a) ALC-ALC (b) ALC-Cholesterol and (c) Cholesterol-Cholesterol lipid pairs. Probability distribution of the splay (S) is shown in blue, the PMF along the splay is shown in green and the parabolic fit to the PMF is shown in red.

#### 6.3 $\kappa$ for MC3 only, ALC only $H_{II}$ phases

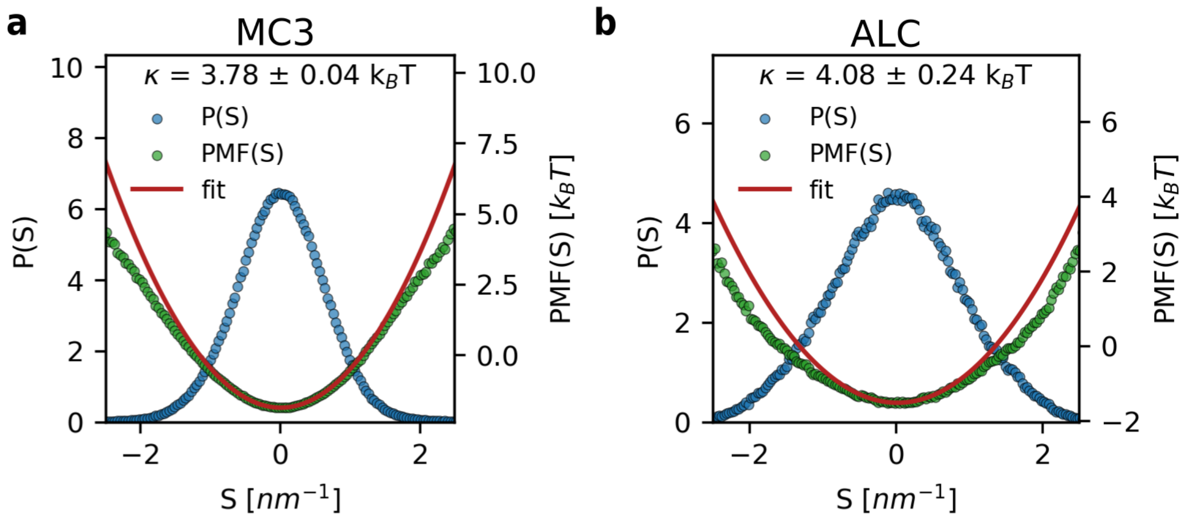

Figure S11: Bending rigidity $\kappa$ obtained from (a) MC3-only $H_{II}$ phases and (b) ALC-only $H_{II}$ phases. Probability distribution of the splay (S) is shown in blue, the PMF along the splay is shown in green and the parabolic fit to the PMF is shown in red.

### 7 Fit of the Laurdan spectroscopy data

#### 7.1 Hysteresis data

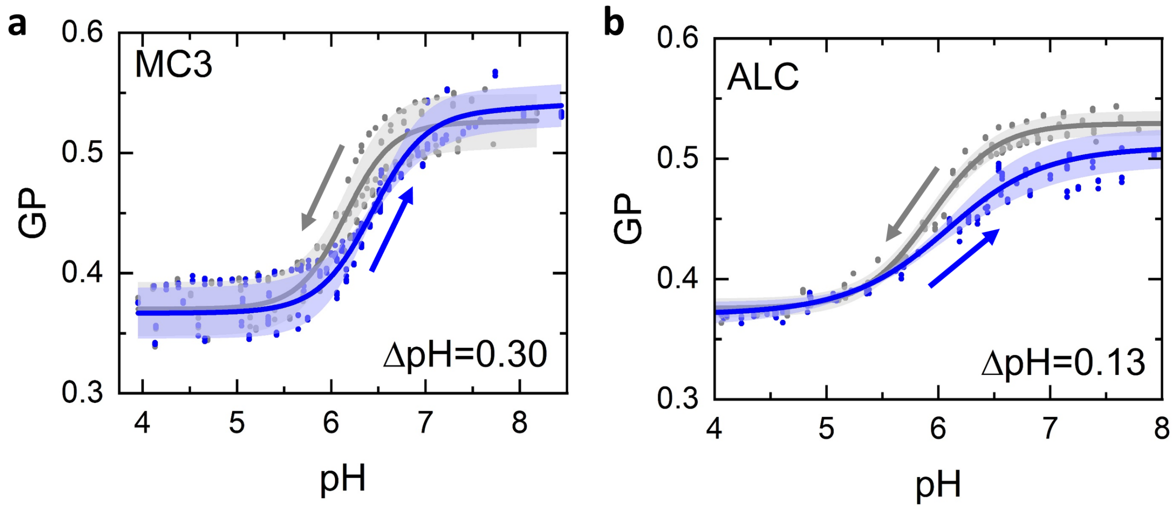

Figure S12: Detection of hysteresis of the phase transition in MC3-containing (a) and ALC-containing (b) LNP measured with the Laurdan fluorescence. The data points correspond to the measurements of 4 LNP replicates, and a triplet measurement at every pH step. Arrows indicate the direction of pH titration.

#### 7.2 Temperature dependence

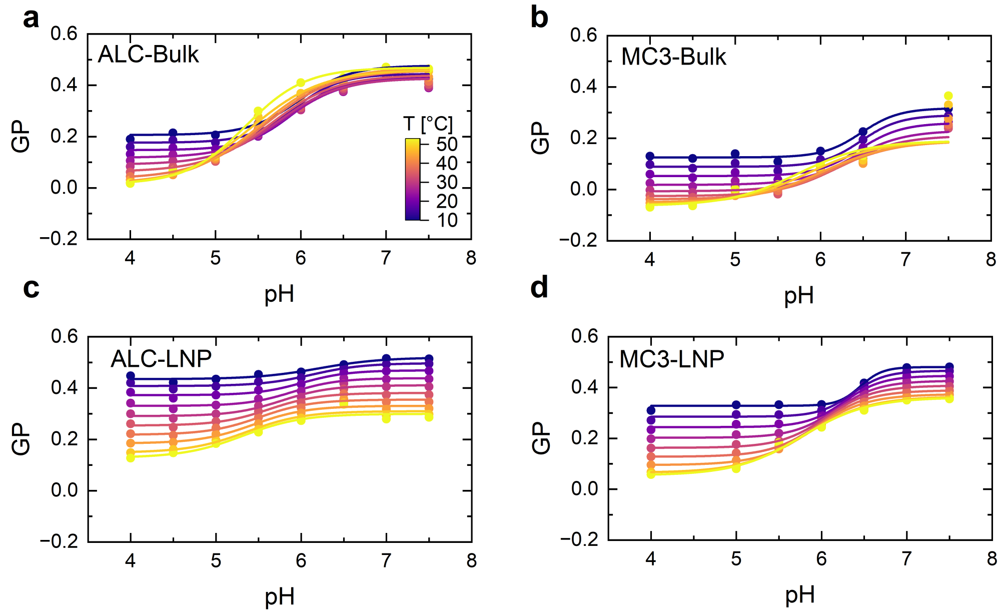

Figure S13: pH- and temperature-dependent changes in GP values of bulk system formulated with ALC-315 (a) and DLin-MC3-DMA (b) and LNPs formulated with ALC-315 (c) and DLin-MC3-DMA (d).
